# Never let me down: new possibilities for lowering serum free cultivation media costs

**DOI:** 10.1101/2022.11.13.516330

**Authors:** Lisa Schenzle, Kristina Egger, Bernhard Spangl, Mohamed Hussein, Atefeh Ebrahimian, Harald Kuehnel, Frederico C. Ferreira, Diana M. C. Marques, Beate Berchtold, Nicole Borth, Aleksandra Fuchs, Harald Pichler

**Author notes:** **Co-first authors** Lisa Schenzle and Kristina Egger are equal contributors to this work and designated as co-first authors. **Correspondence** Aleksandra Fuchs Petersgasse 14/V, Graz 8010, Austria.

## Abstract

Cultivated meat may be a more ethical, environmentally friendly, antibiotic-free meat alternative of the future. As of now, one of the main limiting factors for bringing cultivated meat to the market is the high cost of the cell culture media and their great dependency on serum albumins, production of which is predicted to become a major bottleneck of this industry. Here, using bovine muscle stem cells (BSC) we optimized B8/B9 medium - one of the well-established serum free, fully defined medium compositions available for purchase or for preparation in-house. We show several combinations of the growth factors/myokines/hormones, which were able to substantially increase BSCs proliferation rate, as well as treatment schemes allowing for five to ten times lower concentrations of signaling molecules for the same effect. Additionally, we identified several food grade, low-price medium stabilizers, exhibiting comparable or even superior stabilization of the B8 medium in short-term cultivations, as compared to recombinant human serum albumin (HSA). DoE aided in identifying the best cultivation conditions. Other satellite cells (porcine, chicken and fish) were grown in several final cell culture medium compositions, showing significant cell-line specific differences in performance. Also, the proliferation and yield of CHO cell line, which is relevant e.g. for the production of growth factors, was also successfully increased using our stabilization approach. We conclude that stabilizers tested here can act as versatile low-cost medium additives, partly by prolonging the half-life of growth factors. Thus, we provide an alternative to HSA, enabling up to an overall 73% reduction of medium price.

## Introduction

Conventional meat production and especially beef production forms the tip among the most land and emission intensive food products. Since the world population, and therefore the meat consumption, is predicted to increase by further 73% until 2050, it will not be possible to obtain enough meat in a conventional way, as already around 90% of all agricultural land is used for animal feed^1^. But one of the possible alternatives – cultivated meat – can drastically reduce land use and global warming effects.

Cultivated meat has come a long way from its first introduction in 2013 by Prof. Mark Post, with his first cultivated meat hamburger at € 250,000.-using expensive and ethically questionable Fetal Bovine Serum (FBS) containing medium – a standard those days^2^. Since then, multiple groups have been working on making cultivated meat production feasible and prices low enough to compete with conventional meat on the free market. A lot of effort is still being put into the development of cultivation medium, as over 95% of the production costs are attributed to it^3,4^ due to the very high prices for recombinant growth factors (GFs) and other components used herein – such as serum albumin used in concentrations from 0.8 to 5 g/L^5,6^. In the first attempts, formulations known from work with other types of stem cells, e.g. iPSC cells, were used^7^, but those formulations were not sufficiently effective and still overly expensive. During the last years quite a lot of progress was made in this area. Among other – proprietary – solutions^7,8^, three renown groups have published their serum-free, fully defined medium compositions, which allow very high propagation efficiency, comparable or even superior to FBS-based medium formulations.

For the first optimized medium, Stout *et al.* have taken a further development of the Essential 8 medium - called B8 medium^7^ – as a basis, and supplemented it with 0.8 g/L human serum albumin (HSA) for stabilization. This new medium – called B9 – was published in June 2022^5^, is shown to perform nearly as well as the 20% FBS-containing growth medium on primary bovine satellite cells (BSCs). It’s HSA costs comprised 24.56 USD/L, and overall medium costs based on bulk prices were in the range of USD 46.28 (excluding coating proteins, with 5 µg/L FGF-2) and with HSA making up to over 50% of total cost. Comparing to the Essential 8, B9 medium supports the propagation of satellite cells more effectively, and is approx. six times cheaper than Essential 8 at about 500 USD/L^5,7^.

The other serum-free, fully defined medium was published by Kolkmann *et al.*^6^ in January 2022, where several non-standard GFs and a myokine were introduced (IL-6, IGF1, VEGF, HGF and PDGF-BB – here called PDGF). These exhibit synergistic effects on BSCs proliferation and reach 97% of the efficiency of the 20% FBS-containing growth medium. To produce such effect, Kolkmann used even higher HSA concentration for stabilization – 5 g/L end concentration – for an approximate cost of 163 USD/L only for HSA.

Yet another group has reported very impressive results using a mixture of usual GFs (FGF-2, Insulin, PDGF, HGF), supplemented with the combination of Bovine Serum Albumin (0.075 g/L) and Fetuin (0.6 g/L) – a fetal variant of Serum Albumin, more abundantly present in FBS than Serum Albumin^9^.

At this point, it became apparent that more intricate media were to be developed. These media featured various GFs and signaling molecules that orchestrated the proliferation of stem cells, aiming to replace the extreme complexity associated with FBS. Additionally, this trend was reinforced by using medium stabilization, effectively taking on some of the functions of the extracellular matrix (ECM). Such stabilization simultaneously presents a central proliferation potentiating factor, raising effectiveness of the medium, but is also a cost factor, comparable to or even bigger than the GFs themselves. Moreover, the production volume of recombinant albumin required to replace just 1% of the globally consumed meat would be in the millions of kilograms, surpassing by far the current production volumes of many industrial enzymes^10^.

The most recent paper from Stout *et al.* has addressed this latter issue, demonstrating a successful HSA substitution by in-house produced seed protein isolates, enriched with plant albumins^11^. Extremely low prices of the oilseed protein meals (less than 0.4 USD/kg), which were used as the starting material, do raise hope that the final prices for protein preparations will be rather low. But it is hard to assess this fact properly, as such isolates are currently commercially unavailable, and the proposed isolation method consists of alkali extraction (pH 12.5), isoelectric precipitation (pH 4.5), centrifugation, filtration, and ultrafiltration to concentrate the final protein solutions to 50 mg/mL, as well as the fact that the isolates can only be stably stored at -80°C without the loss of function^11^. Some of these techniques are still quite costly on an industrial scale, though the extraction process, as well as storage conditions can be further optimized.

Here, we suggest several variants of the improved B8 and B9 media as a serum free, fully defined medium composition, which can be prepared in-house. We document combinations of the GFs/myokines/hormones, able to increase the satellite cells’ proliferation rate substantially. We also show that, compared to a single addition of the signaling molecules, two treatments with significantly lower concentrations demonstrate much more pronounced proliferation effect. This finding not only underscores the importance of adjusting the application scheme, as it can potentially lead to a substantial – factor 5x to 10x – cost savings, but also indicates yet again that the stability of GFs is absolutely pivotal to superior performance of culture medium. Additionally, we present several low-priced, food-grade medium stabilizers and their combinations, which exhibit similar stabilization of the B8 medium as compared to recombinant HSA, allowing for its substitution for some cell lines, lowering the price for stabilization to approximately 0.1% of that used in B9 medium. Moreover, we show that combination of HSA with methyl cellulose (MC) exhibits a superior stabilization effect for BSCs, as compared to any stabilizer alone, but not for other species’ satellite cells, emphasizing the species-specificity of cell culture medium optimization regarding ECM mimicking. In sum, our findings allow for a further cost reduction for the propagation phase medium, bringing cultivated meat closer to the market.

## Results

### Single addition of GFs requires much higher concentration for the same effect

GFs are inherently unstable proteins, with some of them having half-lives below 1 h in solution at 37°C^12^. Myokines, which are also known to induce muscle hypertrophy^13,14^, were shown to be rapidly depleted in plasma only 2-4 h after physical activity, and lead to muscle hypertrophy upon repeated physical activity^15^. Considering this, we decided to compare a standard single treatment scheme with a double treatment scheme, which should have simulated a more pro- muscle hypertrophic scenario of a repeated “physical activity” with 2 high picks of components and a low basic level in between. Thus, taking B9 including 0.8 mg/ml HSA^5^, as a basis, we compared the effect of components’ addition on day 1 with adding them on days 1 and 3 on the proliferation of primary bovine satellite cells (BSCs) (please see supplemental figure 1 for characterization of freshly isolated SCs).

As expected, single addition did require much higher concentration for the same or less pronounced effect (**Figure 1**). For example, upon single addition of the recombinant human Hepatocyte Growth Factor (rhHGF), a concentration of 20 ng/mL was necessary for 1.5 fold higher Presto Blue signal (**Figure 1** B), corresponding to a higher cell density, whereas when applied twice, the threshold of 1.5 fold improvement was consistently achieved at rhHGF concentrations of 2.5 ng/mL (**Figure 1** A). The same effects, but even more pronouncedly, were observed e.g. upon Estrogen (17β-Estradiol) addition (**Figure 1 C-D**). This emphasizes the significance of adapting the treatment scheme to conditions that are more physiologically relevant, as it has the potential to result in substantial cost savings, possibly up to a factor of 5x or 10x, on the most expensive medium components.

**Figure 1:**
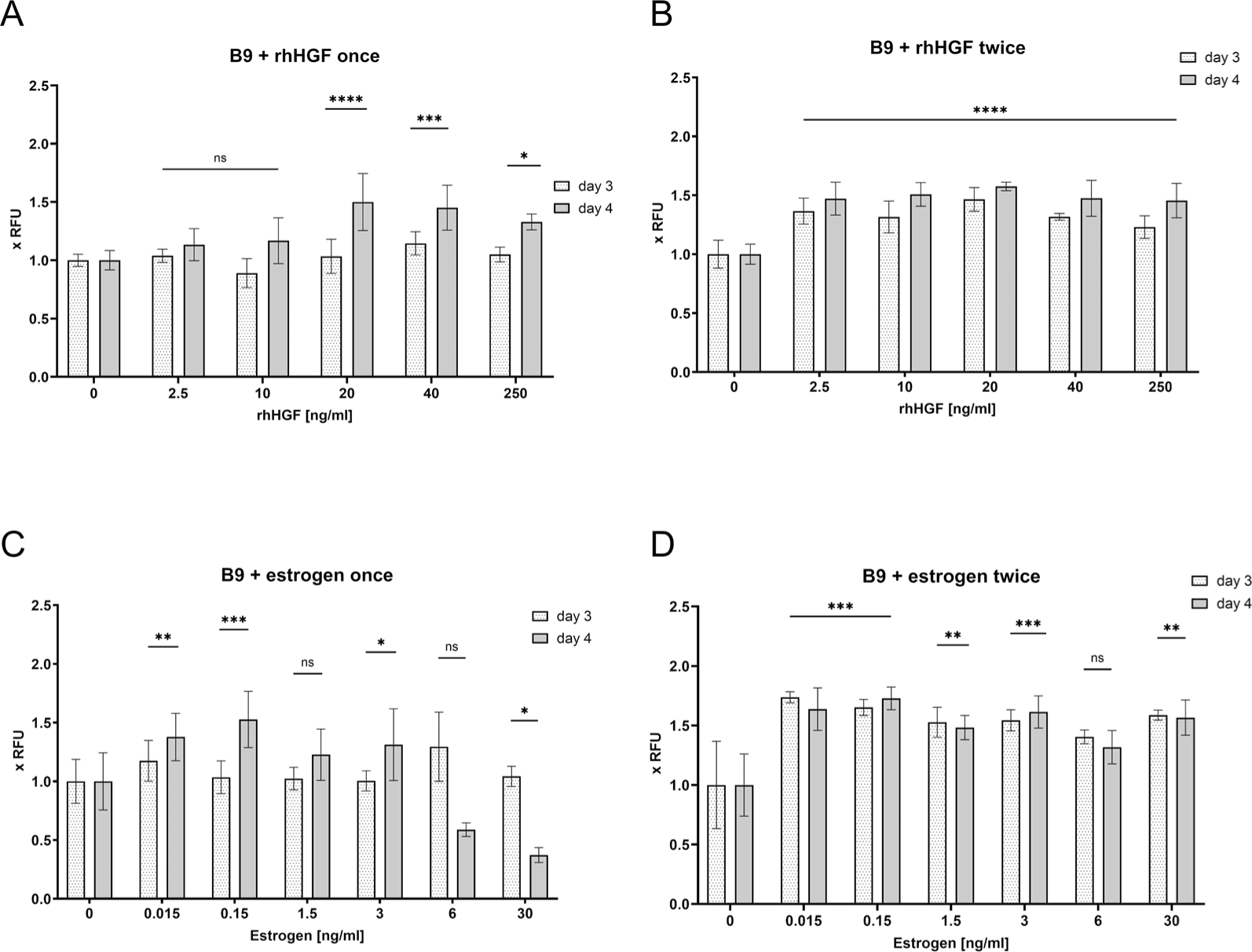
Comparison of single and double addition of components. BSCs were treated either once on day 1 or twice on days 1 and 3 with indicated concentrations of the rhHGF **(A and B)** and estrogen **(C and D)**. Presto Blue assay was performed on days 3 and 4, and normalized to vehicle treated control (0), which contained DPBS + 0.8 mg/mL HSA (B9). n=6 biological replicates for **A** and **C**, n=3 for **B** and **D**; statistical significance was calculated by one-way ANOVA combined with Dunnett test for day 4, comparing all samples with vehicle treated control (0), and is indicated by asterisks, which are p < 0.05 (*), p <0.01 (**), p < 0.001 (***).

Regarding most other potential proliferation inducers (see supplemental table 1), the effect of a double treatment was more pronounced than with a single treatment (data not shown). Therefore, in all subsequent experiments, we utilized a double treatment of the components.

### Screening single medium components

The addition of individual components to the serum-free medium is the traditional method of medium optimization. This approach aims to minimize the number of components to simplify media preparation under laboratory conditions, though it is a challenge to reach the same proliferation efficiency as it is possible with complex FBS-containing media, where molecular surrounding consists of hundreds of active molecules in all ranges of concentrations^16^.

To this end, we have tried out addition of single components to B9 medium, and combinations of the resulting hits. Most of the tested components elicited a pronounced positive effect on the proliferation of bovine satellite cells, when applied twice. In total, 19 components were chosen based on literature (see supplemental table 1 for the whole list and sourcing), from which the most prominent hits where rhHGF, Estrogen (17β-Estradiol), rbIL-6, rmWnt3a, and moderately good hits – rhIL1α, rhPDGF-BB, Testosterone (5α-Dihydrotestosterone), based on the Presto Blue assay (**Figure 2**). See supplemental figure 2 for the proliferation timelines of the rest of individual components – rhFollistatin, rhCHI3L1, rhWnt-5b, rhIGF1, rhIL-4, rhIL-13, rhIL-15, rhLIF, rhIFN-γ, rhTNF-α, rhGASP-1, EGF.

**Figure 2:**
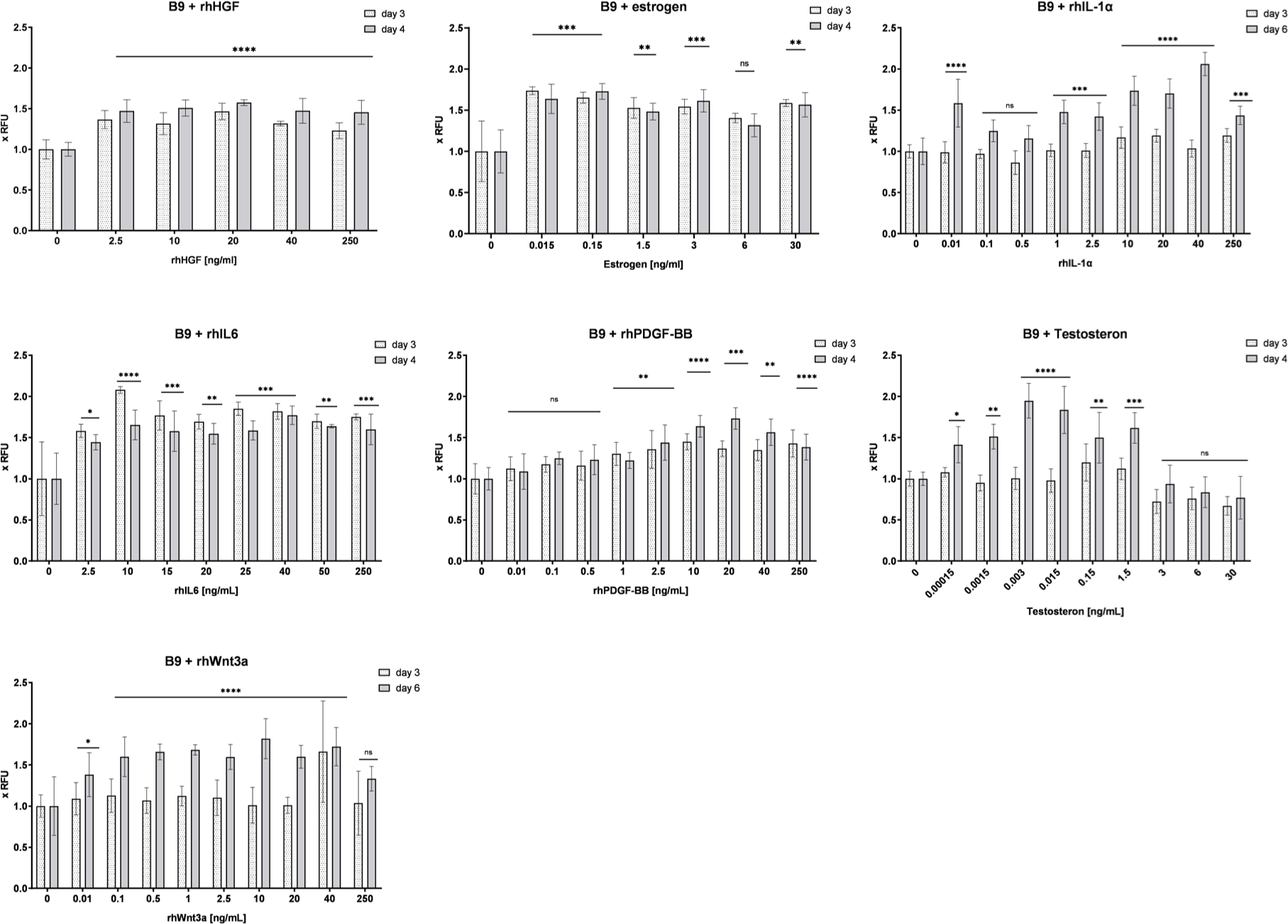
Screening single medium components for induction of proliferation. BSCs were treated on days 1 and 3 with indicated concentrations of components, and Presto Blue assay was performed on days 3 and 4, or 3 and 6. Obtained values were normalized to vehicle treated control (0), which contained DPBS + 0.8 mg/mL HSA (B9). n=3 (rhHGF, IL-6, Estrogen) and n=6 (rhPDGF-BB, testosterone, rhWnt3a, rhIL-1α) biological replicates, experiment repeated at least twice; statistical significance was calculated by one-way ANOVA combined with Dunnett test for day 4 or 6, comparing all samples with vehicle treated control (0), and is indicated by asterisks, which are p < 0.05 (*), p <0.01 (**), p < 0.001 (***), p < 0.0001 (****).

It has to be mentioned that MC at high concentrations used for preparation of stock solutions – at 5 g/L – is quite difficult to handle due to its high viscosity, which means that the two highest concentrations (0.9 and 0.45 g/L) could be less precise.

Presto Blue is a viability assay representing cells’ metabolic activity and thus indirect in its nature^17^. To ensure that the data produced by this assay is relevant under the conditions where the added proliferation inducing compounds could cause a change in the cells’ metabolic activity, a direct assay was used to assess the quantity of DNA per well, and thus cell density more accurately at confluence and below – Hoechst 33258 assay. This assay can be used for cell densities up to 100.000 cells/cm^2^. Such combination of two assays was previously reported to function very reliably also in a high-throughput format^6^. We thus conducted Hoechst assay on the last day of respective experiment, which mostly confirmed our earlier findings (see supplemental figure 3), except that in the experiments lasting 8 days (rhPDGF-BB, testosterone, rhWnt3a, rhIL-1α), the positive effect of component addition was flattened out as all samples reached the confluence.

### Screening combinations of the resulting hits

For the screening of combinations, the lowest concentration which had had a positive effect in the screening was chosen, to avoid possible adverse effects sometimes seen at high concentrations, and to keep the medium price as low as possible. The screening was performed as above – Presto Blue assays were performed on days 3 and 4, and additionally on days 6 or 7. Presto Blue assays were less reliable at higher cell densities around confluence, reached on days 6-8 (approx. 100,000 cells/cm^2^). Thus, the significance of the observed effects was statistically evaluated by one-way ANOVA combined with Dunnett test calculated for day 4 or at the latest day 6. Additionally, the data was confirmed by Hoechst assay at the end of the same screening round (data not shown).

We could identify following hits - see **Table 1** (see supplemental figure 4 for the proliferation timelines of individual combinations).

**Table 1:** Summary of the screening analysis of the single components and simple combinations based on Presto Blue screening. Darker marked components elicit more prominent effect on proliferation rate of BSCs. Unmarked components elicited least prominent effects (under 1.5 x RFU). N.A. – not analysed. * - unstable effect.

| Component | B9 + 1 component added once | B9 + 1 component added twice | B9 + 2 components added twice<br>(concentration = 2.5 ng/mL<br>Estrogen (EST) = 0.15 ng/mL) |  |  |
| --- | --- | --- | --- | --- | --- |
|  |  |  | +rhHGF | +IL-6 | + EST |
| Estrogen | from 0.15 ng/mL | from 0.015 ng/mL |  |  |  |
| rhHGF | from 20 ng/mL | from 2.5 ng/mL |  |  |  |
| rhIL-6 | from 40 ng/mL | from 10 ng/mL |  |  |  |
| rhWnt-3a | N.A. | from 0.5 ng/mL |  |  |  |
| rhPDGF-BB | N.A. | from 10 ng/mL |  |  |  |
| rhFollistatin | N.A. | from 10 ng/mL |  |  |  |
| rhCHI3L1 | from 0,1 ng/ml* |  |  |  |  |
| rhIGF1 | N.A. | from 2.5 ng/mL |  |  |  |
| rhWnt-5b | N.A. | from 10 ng/mL* |  |  |  |
| rhIL1α | N.A. | from 10 ng/mL |  |  |  |
| Testosterone | N.A. | from 0.015 ng/mL |  |  | 1:5 |

From 19 components tested, the best, most consistent results upon double addition of a single component to B9 medium were shown for rhHGF (Abcam #ab245957), second best - 17β-Estradiol (Sigma #E2758-250MG), third best – rbIL-6 (Biorad #PBP021). Best combinations of two components were rhHGF + rhPDGF-BB, second best – rhHGF + rmWnt3a and and IL-6 + rhPDGF-BB, third best – rhHGF + rhWnt-5b.

Further on, to determine the most potent combination of our hits, the best two-component combinations - rhHGF + rhPDGF-BB – was gradually expanded to include additional best two-component hits – IL-6, Estrogen and Wnt-5b+Wnt3a. The combination of Wnt-5b+Wnt3a was previously published to promote YAP/TAZ activation via the alternative Wnt signaling pathway^18^. Also, both were among the best hits in combination with rhHGF (**Table 1**) and had a positive effect when combined without rhHGF (data now shown). They were thus included into the final screening together.

As shown in **Figure 3**, Presto Blue assay data suggests that in B9 medium on day 4 the prominent effect was demonstrated by IL-6 addition to rhHGF + rhPDGF, though it could further be slightly improved in the combination of rhHGF + rhPDGF + IL-6 + Wnt3a + Wnt5b, as well as in the combination rhHGF + rhPDGF + IL-6 + Wnt3a + Wnt5b + Estrogen, but less so. It was surprising to see no or a negative effect from addition of Estrogen to rhHGF + rhPDGF + IL-6, because it showed significant improvement in several combinations – especially with rhHGF and rhPDGF (see Suppl. Fig. 2). Estrogen concentrations below 5 ng/mL were further on tested in the same setup, but without any significant advantage upon its addition (data not shown).

**Figure 3:**
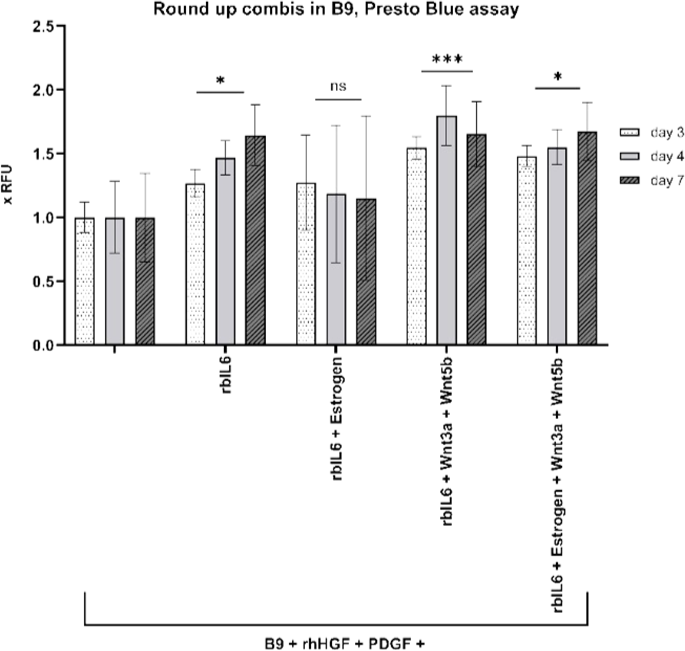
Final combinations of proliferation inducing hits in B9 medium. BSCs were treated with 5 ng/mL of each component on days 1 and 3, Presto Blue assay was performed on days 3, 4, and 7. Obtained values were normalized to BSCs treated with rhHGF + rhPDGF. Data for n=6 biological replicates is presented; statistical significance was calculated by one-way ANOVA combined with Dunnett test for day 4 (Presto Blue), comparing all samples to BSCs treated with rhHGF + rhPDGF, and is indicated by asterisks, which are p < 0.05 (*), p <0.01 (**), p < 0.001 (***), p < 0.0001 (****).

### Non-specific stabilization of cultured medium components

Albumins are very widely used for non-specific stabilization of biologically active proteins^19–21^. But they are far from being the only known protein stabilizers – salts, sugars, amino acids, fatty acids and hydrogels are also used for this purpose, some of them known under the term “chemical chaperones”^12,22,23^. Also, different GFs can require various stabilizers at specific concentrations for a significant effect^21,24^. However, sustainable stabilization properties for longer storage are often reached at extremely high, non-physiological concentrations and could be causing serious adverse effects^25,26^.

Thus, the effect of two alternatives to recombinant HSA were investigated – methyl cellulose (MC) and DL-alanine (ALA), which are known for their non-specific stabilization effects of FGF-2^24^ (which is one of the components of B8 and B9 media), are extremely cheap in comparison to HSA (see **Table 2**), and are both food components – MC is an approved food emulsifier also known as E461, and L-ALA is a natural food compound. Instead of L-ALA we used racemate to follow-up on the effects published by Benington *et al*.^24^ We hypothesized that such non-specific stabilizers can achieve additional positive effect, regardless of the GFs combination ultimately used.

**Table 2:** Cost comparison and sourcing of different stabilizing agents used in this study.

| Compound | Article Nr | Cost | Costs/L medium |
| --- | --- | --- | --- |
| rhAlbumin | Oryzogen OsrHSA | € 25.-/kg | 0.8 g/L → 26.12 €/L |
| DL-Alanine, ≥99%, FCC, FG | Sigma W381810-10KG-K | € 61.2/kg | 0.4495 g/L (5 mM) → 0.0275 €/L |
| Methyl cellulose viscosity: 4,000 cP | Sigma M0512-1KG | € 328.-/kg | 0.1125 g/L → 0.037 €/L |
| Starch (corn) | Merck # S4126-5KG (5 kg) | € 42.8/kg | 0.4 g/L → 0.017 €/L |
| Starch (rice) | Merck # S7260-1KG (1 kg) | € 76.50 /kg | - |
| D-Sorbitol | Roth # 6213.7 (25 kg) | € 13.56/kg | 0.4 g/L → 0.005 €/L |
| D-Mannitol | Roth # 4175.2 (5 kg) | € 83.8/kg | 0.1 g/L → 0.008 €/L |
| Inulin | Thermo # 457105000 | € 1,142.-/kg | - |
| Locust bean gum | Merck # G0753-1KG | € 163.-/kg | - |
| Cyanoflan | Gift from Dr. Rita Mota<sup>27,28</sup> | Costs unknown | 0.8 g/L → Costs unknown |

Stabilizing activity of methyl cellulose (MC) and alanine (ALA) towards less stable B8/B9 components (Insulin, FGF2, TGFβ3, NRG1)^5^ was investigated in standard cell proliferation assays on BSCs in either B8 (without 800 mg/L HSA) or B9 (with 800 mg/L HSA) media. Several concentrations were applied around 0.5 g/L MC and 20 mM ALA, known to have high stabilizing potential for FGF-2^24^. The combinations of all three stabilizers (HSA, MC and ALA) were also tested, as we reasoned it would be mimicking the very generalized content of ECM, consisting mostly of water, proteins, amino acids and polysaccharides^29^.

As evident from **Figure 4** A, in short term proliferation, both MC and ALA alone and in combination could stabilize B8 approximately as well as HSA at the lower concentrations used: MC at 0.1125 g/L, ALA at 10 mM and 5 mM (trend supported by Hoechst data, **Figure 4** C), with MC alone performing slightly better than ALA alone. But even more interesting was the superior stabilization of B9 medium upon addition of MC or MC + ALA at the lower applied concentrations (**Figure 4** B), reaching significance in Hoechst assay (**Figure 4** D). Also, lower concentrations of MC at 0.1125 g/L and ALA at 5 mM were examined, but with no further improvement of the proliferation rates (data not shown).

**Figure 4:**
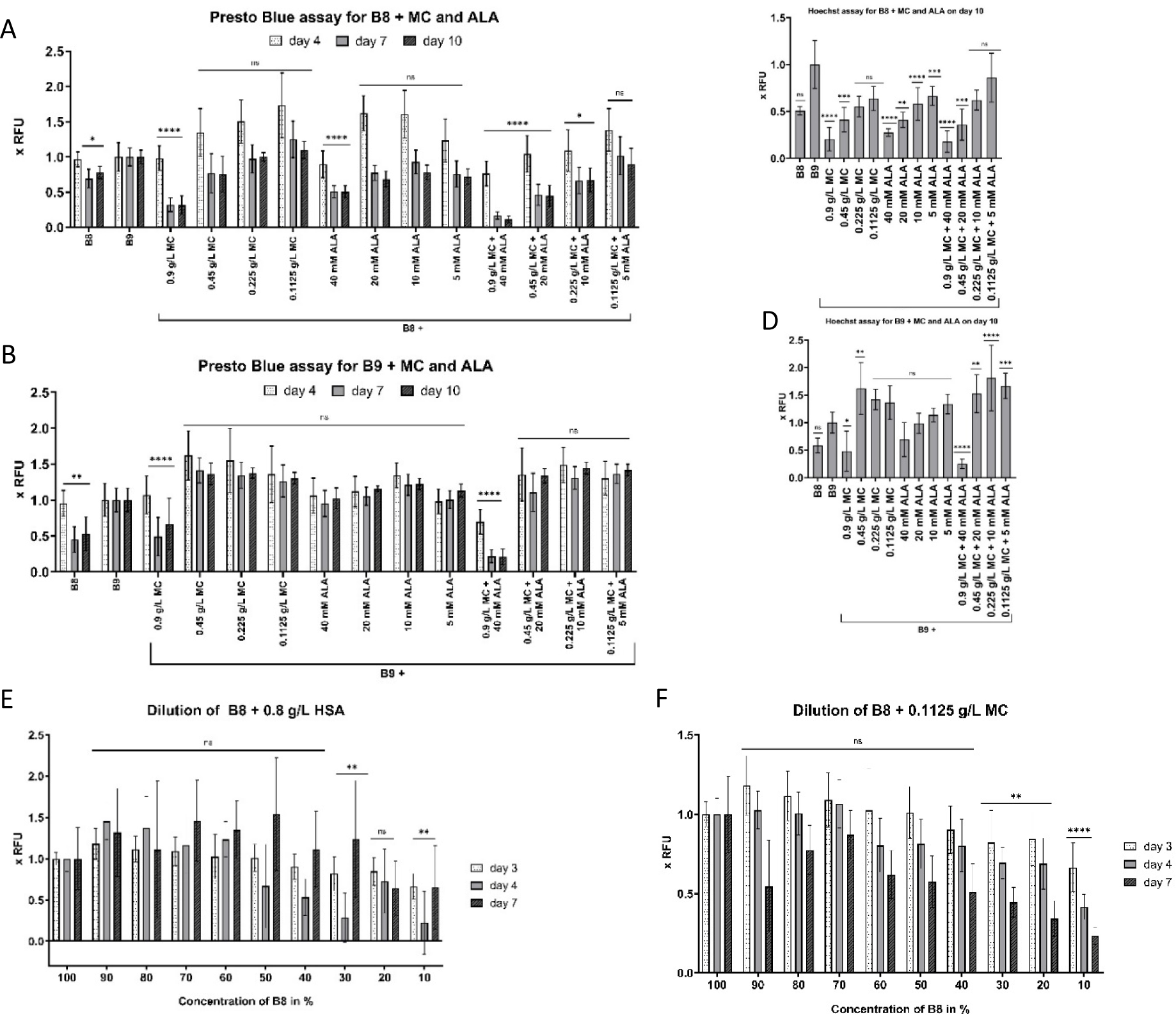
Non-specific stabilization of cultured medium components. 2000 BSCs/cm^2^ were seeded on day 0 in BSC-GM, and changed on day 1 to either B8 or B9 medium, with methyl cellulose (MC) and/or alanine (ALA) added to indicated end-concentrations with every medium exchange. Presto Blue assay was performed on indicated days **(A, B, E, F)**, and Hoechst assay – on day 10 **(C and D)**. Obtained values were normalized to BSCs in B9 **(A-D)** or to the values of basic (100%) B8 concentration **(E-F)**. n=6 biological replicates and repeated at least twice; statistical significance was calculated by one-way ANOVA combined with Dunnett test for day 4 (Presto Blue) or for day 10 (Hoechst), comparing all samples to B9 **(A-D)** or to the values of basic (100%) B8 concentration **(E-F)**, and is indicated by asterisks, which are p < 0.05 (*), p <0.01 (**), p < 0.001 (***), p < 0.0001 (****).

It must be mentioned that MC at high concentrations used for preparation of stock solutions – at 5 g/L – is quite difficult to handle due to its high viscosity, which means that the two highest concentrations (0.9 and 0.45 g/L) could be less precise.

All in all, the most prominent stabilization of B8 – on the level of 0.8 g/L HSA – was achieved by MC (0.1125 g/L) or by the combination MC + ALA (0.1125 g/L + 5 mM respectively). Most prominent stabilization of B9 – at least 1.5x better than B9 -with a triple combination of HSA (0.8 g/L), MC (0.45-0.1125 g/L) and ALA (20-5 mM). As Stout *et al.* have shown that HSA can be substituted by plant seed protein isolates^30^, this could make this triple combination commercially feasible.

Considering that the chosen ALA compound – racemate as was published by Benington *et al.*^24^ – is not a food grade component because it contains D-alanine, we further tested L-ALA instead of the racemate in combination with MC in B8 and B9 media and found the same but not more prominent effects (data not shown). As L-ALA is very cheap it could be reasonably expected to potentiate some GFs in other settings.

In the further effort to lower the medium costs, the effect of lowering B8 content containing Insulin, Transferrin, FGF2-G3, TGFβ3, NRG1 as active components, with HSA or MC as stabilizers was tested. Therefore, the viability screen as previously described was performed with a B8 proportion in DMEM-F12 gradually lowered, whereas concentrations of stabilizers stayed constant. Statistical analysis shows that the dilution of B8 in DMEM-F12 stabilized by either 0.1125 g/L MC or by 0.8 g/L HSA does not lead to a significant drop in viability on day 4 until the concentration of B8 is below 40% (**Figure 4 E-F**). Also, the reduction by the first 30% (MC) - 40% (HSA) did not have any apparent negative effects.

### Stabilizers mechanism of action

First, we have investigated whether the degradation of our most important GFs is actually influenced by the addition of stabilizers MC and L-ALA at 37°C.

rhHGF half-life of 1 h in DPBS at 37°C was slightly higher than that of rhPDGF with 45 min (**Figure *5*** A and B), and the degradation of both rhHGF and rhPDGF was partly circumvented by the addition of MC at 0.1 g/L concentration. L-ALA addition was slightly advantageous in case of rhHGF, but with a more pronounced effect on rhPDGF stability in the first 2 h, which underlines the specificity of stabilizer combinations for each GF.

**Figure 5:**
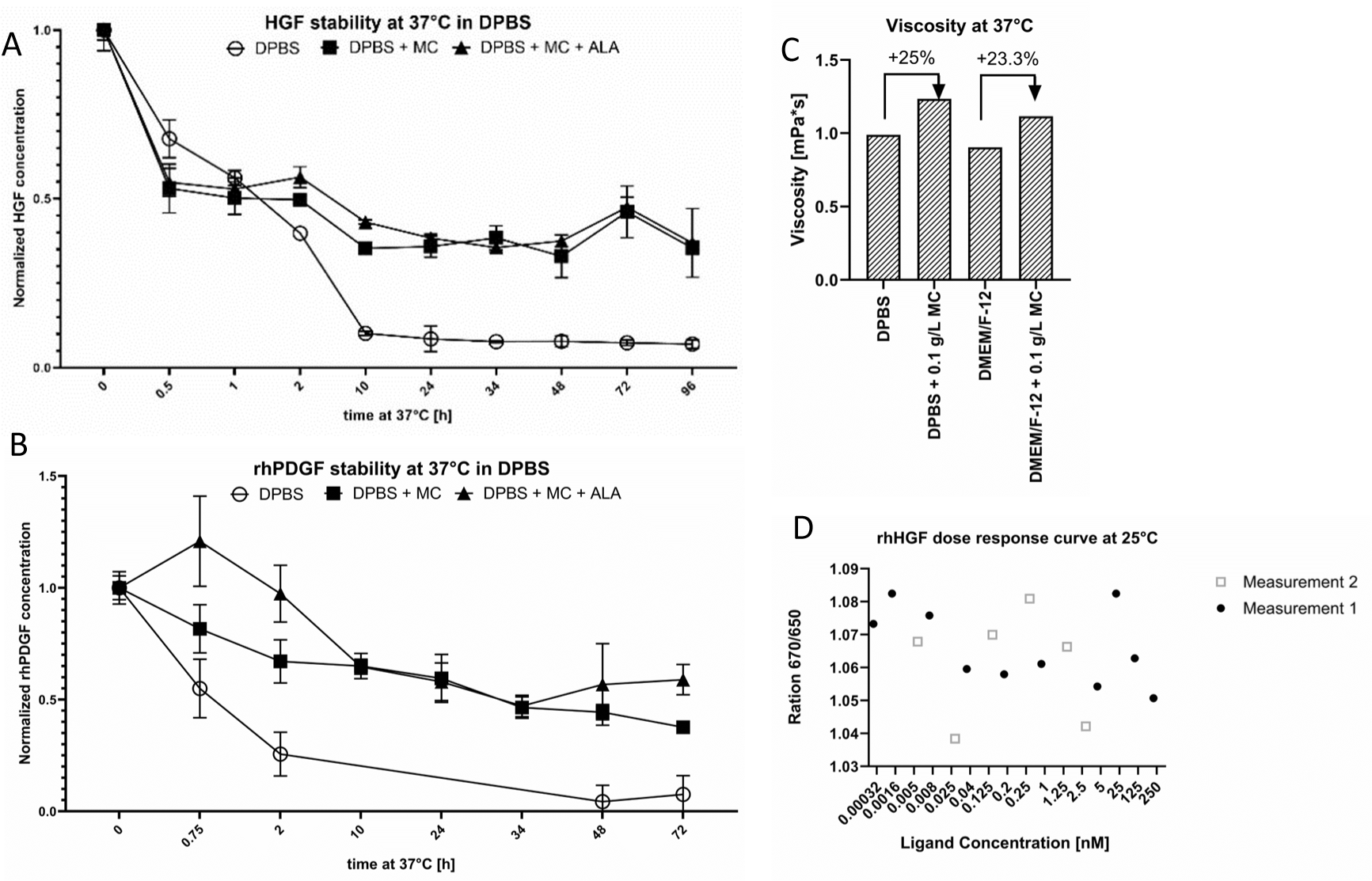
Stabilizers mechanism of action. Stability of rhHGF and rhPDGF at 37°C in DPBS was measured by ELISA following manufacture’s manuals **(A and B)**. GFs in DPBS were stabilized either with 0.1 g/L MC or 5 mM L-ALA. DPBS without stabilizers was used as control. All samples were aliquoted on ice, incubated for the indicated time at 37°C, then frozen (including time point 0), and ELISA was performed after freezing the last sample. **(C)** Viscosity of the buffers and media with/without 0.1 g/L MC was measured using Anton Paar Rheometer MCR 502 following instruction manual and calculated as a function of [shear stress, Pa]/[shear rate, 1/s] over 100 measuring points. **(D)** Change in rhHGF conformation through direct interaction with MC as ligand was not detected. NanoTemper Thermophoresis assay was performed at 25°C with 25nM HIS-tagged rhHGF labeled with 25 nM RED-Tris-NTA dye via binding to its His-tag.

We further compared the viscosity of the solutions with 0.1 g/L methyl cellulose, as viscosity is one of its features, that, at 10x to 50x times higher concentrations is known to contribute to a higher cell survival by acting as an inert adhesion-promoting matrix, possibly through the induction of a higher actin density and remodeling of cytoskeleton^31^. But on the other hand, it could be disadvantageous for quickly proliferating cells, as shown in **Figure 4** A-B. At the concentration of 0.1 g/L, the viscosity attributed to MC was detectibly around 23-25% higher (**Figure *5*** C), and could possibly play a role in higher proliferation by promoting cell adhesion.

Osmolarity can also play a big role in cellular processes. Methyl cellulose was utilized at concentrations of up to 0.09%. Consequently, any fluctuations in osmolarity resulting from the addition of methyl cellulose on our estimation are limited to a maximum of 15 μOsm. This variation is negligible when compared to the osmolarity range of basal media, which is approximately 3,000 times higher, at around 300 mOsm.

Another aspect of how higher concentrations of stabilizers function is by reducing the protein’s free energy through interactions such as ionic, electrostatic, and hydrophobic interactions^24,32^. Specific binding of ligands tends to stabilize a variety of proteins – from enzymes to GFs^33–36^, thus we investigated whether the stabilizing effect of methyl cellulose on GFs is due to such high energy interactions. For that we investigated the direct interaction between rhHGF and MC, as it has demonstrated a more pronounced effect on rhHGF stability. We used NanoTemper Thermophoresis assay to measure the change in GF conformation upon binding to MC. But, as shown in **Figure *5*** D, there seem to be no apparent trend in conformational change of rhHGF in the wide range of MC concentrations (0.00032 nM to 250 nM).

### Stabilization of the best proliferation inducing combinations

After identifying MC as the better performing stabilizing agent for BSCs in B8 and B9 media, MC was applied for stabilization of our best performing proliferation inducing combinations. It is recognized that various stabilizers may exert differing degrees of influence on different growth factors^21,24^. Thus, we compared the stabilization effect of HSA to that of MC for the best component combinations in B8 and in B9 media.

In short term experiments, stabilization of any combination with MC has a significant positive effect on proliferation, as compared to stabilization with HSA (**Figure 6**). The observed effect was more pronounced than during the MC stabilization of B8 without additional growth factors (**Figure 6** compared to **Figure 4** A, C), which could be an indication of MC’s high stabilization capacity towards rhHGF and/or rhPDGF, as compared to stabilization capacity towards Insulin, FGF2, TGFβ3, NRG1^5^. This assumption is supported by the fact that the demonstrated effect was concentration-dependent, and declined with lower concentrations (e.g. 2.5 ng/ml), though multiple components at high concentrations (10 ng/ml) were also detrimental (**Figure 6** A and D). As in the screening for best final combination in B9 medium (**Figure 3**), best combinations were B9 + rhHGF + PDGF + IL-6 and rhHGF + PDGF + IL-6 + Wnt3a + Wnt5b; in B8 stabilized by MC best combinations were B8 + rhHGF + PDGF, and B8 + rhHGF + PDGF + IL-6 + Wnt3a + Wnt5b.

**Figure 6:**
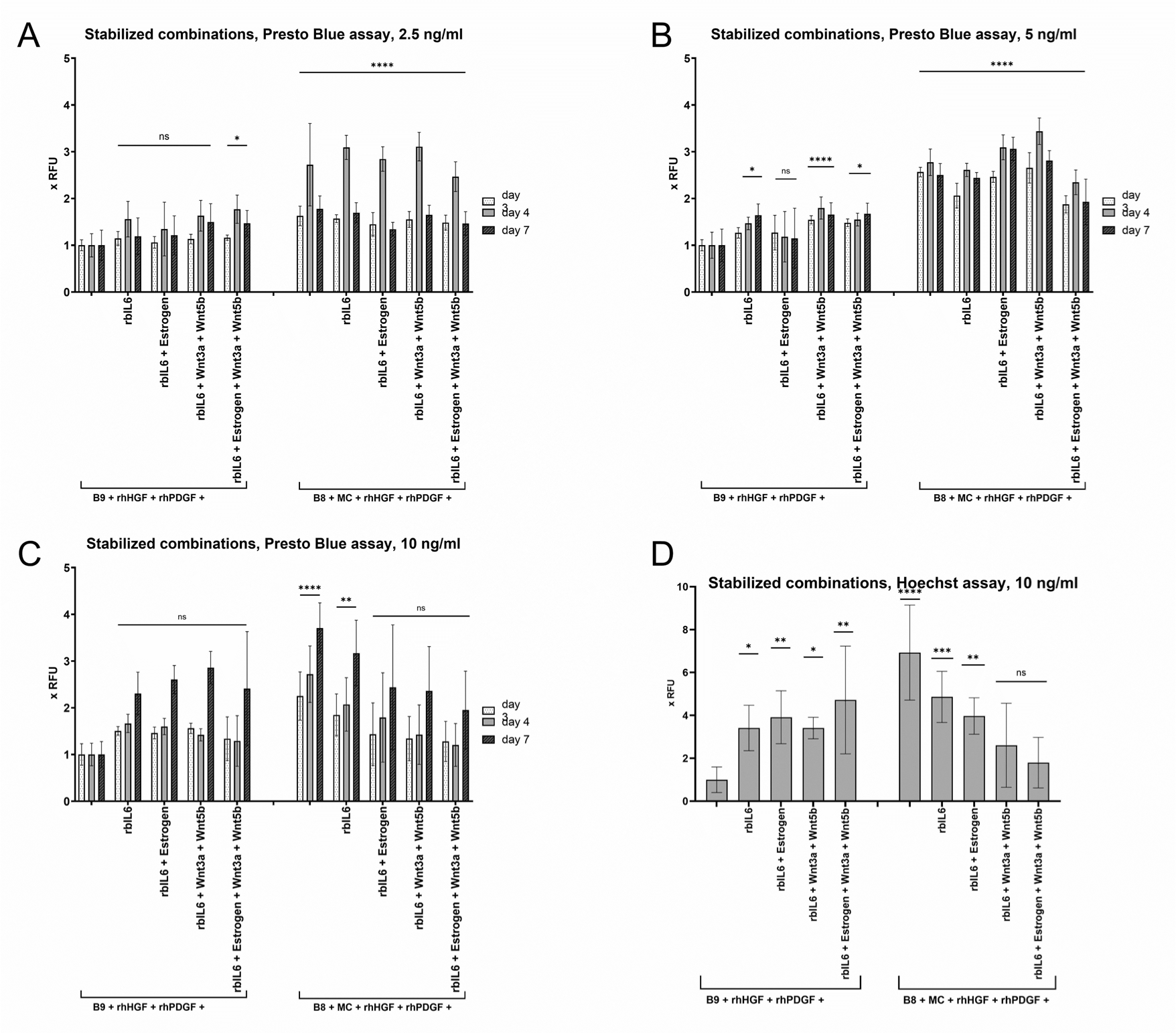
Comparison of stabilization effects of HSA and MC for best proliferation inducing combinations in B8/B9 medium. 2000 BSCs/cm^2^ were seeded on day 0 in BSC-GM, and changed on day 1 to B8 medium, stabilized with either 0.8 g/L human serum albumin (HSA) (B9) or 0.1125 g/L methyl cellulose (MC). Best proliferation inducing combinations were added on day 1 and day 3 to an end concentration of 2.5, 5 or 10 ng/ml, whereas stabilizers were added each time with medium exchange. Presto Blue assay was performed on days 3, 4, and 7 **(A-C)**, and Hoechst assay on day 7 **(D)**. Obtained values were normalized to B9 + rhHGF + rhPDGF. n=6 biological replicates and repeated at least twice; statistical significance was calculated by one-way ANOVA combined with Dunnett test for day 4 (Presto Blue) or for day 7 (Hoechst), comparing all samples with B9 + rhHGF + rhPDGF, and is indicated by asterisks, which are p < 0.05 (*), p <0.01 (**), p < 0.001 (***), p < 0.0001 (****).

### Screening for further alternative stabilizers

Lately, there has been criticism regarding MC inclusion in food items due to its synthetic nature^37^, as it is manufactured by heating cellulose with a caustic solution (such as sodium hydroxide) and subsequently treating it with methyl chloride^38^. We thus explored other possible stabilizers, which would not raise questions about their inclusion into the final food product – e.g. food-grade polysaccharides, with certain structural similarity to methyl cellulose. We also took into account the option of using substances that are metabolically inert.

We could identify 3 further polysaccharides with a similar or slightly better effect on proliferation rates, as compared to B9 or B8+MC (see **Figure 7** for Presto Blue and supplemental figure 5 for Hoechst data) – sorbitol, starch from corn, and locust bean gum. The latter was not included into further experiments due to extremely high viscosity even at low concentrations and tendency to produce aggregates, which led to inconsistent handling and ill reproducibility.

**Figure 7:**
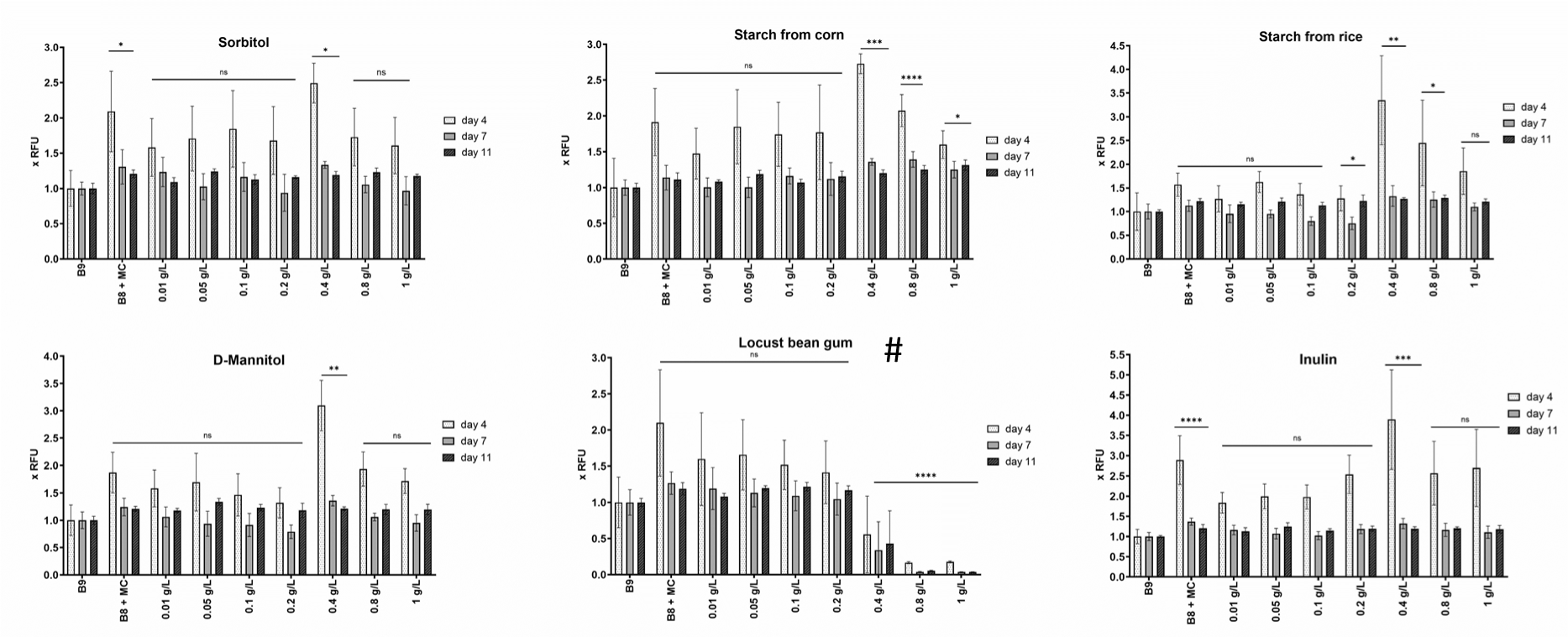
Non-specific stabilization of cultured medium components by new potential stabilizers. 2000 BSCs/cm^2^ were seeded on day 0 in BSC-GM, and changed on day 1 to the designated medium. B8 with 0.1 g/L methyl cellulose (MC) and B9 medium were used as controls. Stabilizers were added to indicated end-concentrations with every medium exchange. Presto Blue assay was performed on indicated days. Obtained values were normalized to BSCs in B9. #LBG concentrations probably not accurate because of aggregation during storage. n=6 biological replicates and repeated at least twice; statistical significance was calculated by one-way ANOVA combined with Dunnett test for day 7, comparing all samples to 100% B9, and is indicated by asterisks, which are p < 0.05 (*), p <0.01 (**), p < 0.001 (***), p < 0.0001 (****).

### Optimization of stabilizer combinations

We employed the design of experiments (DoE) methodology to uncover potential synergies among combinations of stabilizers and to fine-tune their concentrations. Space-filling design^39^ with 6 replicates was applied in a short-term proliferation experiment with 6 stabilizers in concentration ranges listed in **Table 3**, based on data shown in **Figure 7**. Combinations of 3 components in maximum concentrations were excluded from the design to avoid possible adverse effects of higher viscosity. Proliferation of BSCs was measured with Presto Blue assay and confirmed with Hoechst assay, and Presto Blue values, normalized to B9, were analyzed using R^40^ with fitting of linear and linear mixed effects models of second order by stepwise forward regression. Additionally, raw data and background corrected data was analyzed in the same way, yielding very similar outputs (data not shown). The experiment was repeated twice.

**Table 3:** Concentration ranges of stabilizers in DoE setup.

| Starch from corn (STC), HSA, ng/ml | MC, ng/ml | Sorbitol (SO), Mannitol (MA), Inulin (IN), ng/ml |
| --- | --- | --- |
| 0 | 0 | 0 |
| 0.005 | 0.002 | 0.01 |
| 0.01 | 0.01 | 0.05 |
| 0.05 | 0.02 | 0.1 |
| 0.1 | 0.04 | 0.2 |
| 0.2 | 0.08 | 0.4 |
| 0.4 | 0.16 | 0.8 |
| 0.8 | 0.2 | 1 |

The cell density was chosen as an aim of optimization. Influence of stabilizers on proliferation and their reciprocal effects where analysed on single days (dayExp 4, 6 and 8) separately, as well as combined (Table 4). From the effects of single components, MC was the only component which effect reached significance on the two of three single days (days 4 and 6), whereas starch from corn (STC^2, the quadratic term of STC) was significantly advantageous only on day 1. Also, only MC and STC^2 effects reached significance in the analysis of all three days.

We have also analyzed cell density as a function of possible interplay of multiple components, and most significant effects of the model, predicting the proliferative effects throughout days 4 to 8, are presented on **Figure 8**. In this setup, the most advantageous conditions were in higher concentrations of single components, though several components (STC, MC, SO) have demonstrated additive positive effects. From the combinations of 6 tested components, only MC and SO have shown somewhat synergistic effect on day 8 (MC:SO), but without reaching significance – p=0.19 (**Table 4**). Interestingly, combination MC:HSA, which seemed more advantageous than MC alone in short term (**Figure 4** A-B), was shown here rather adversary, with the same trend of favorable effects in cases of either HSA of MC alone.

**Figure 8:**
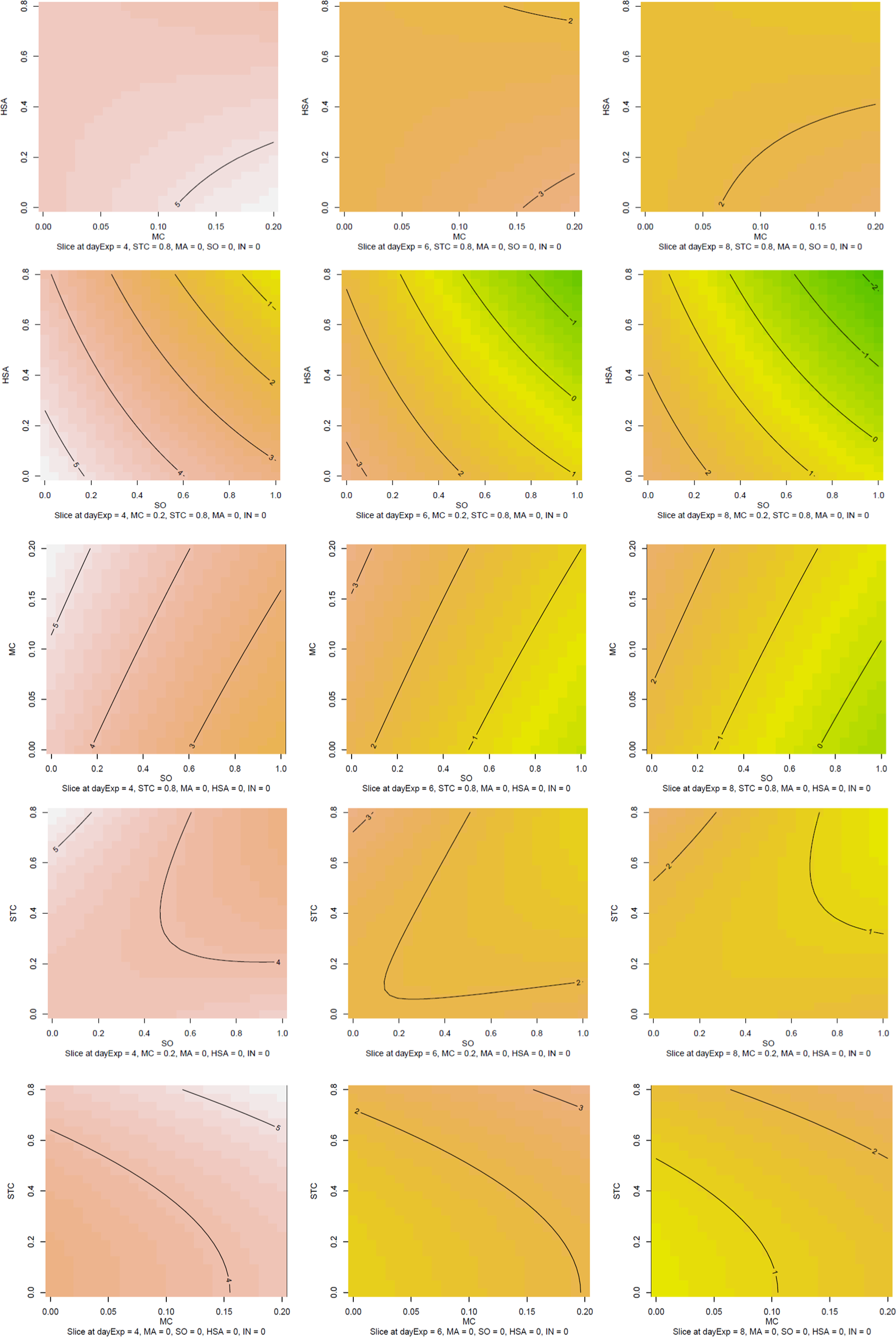
Response surface contour plot predicting the cell density as a function of stabilizer combinations for days 4-8. Optimization of stabilizers concentrations was performed, where starch from corn (STC), recombinant human serum albumin (HSA), methyl cellulose (MC), sorbitol (SO), mannitol (MA) and Inulin (IN) were applied in combinations at concentrations listed in Table 3. The statistics software R^40^ was used for the analysis of the normalized to B9 values.

**Table 4:**
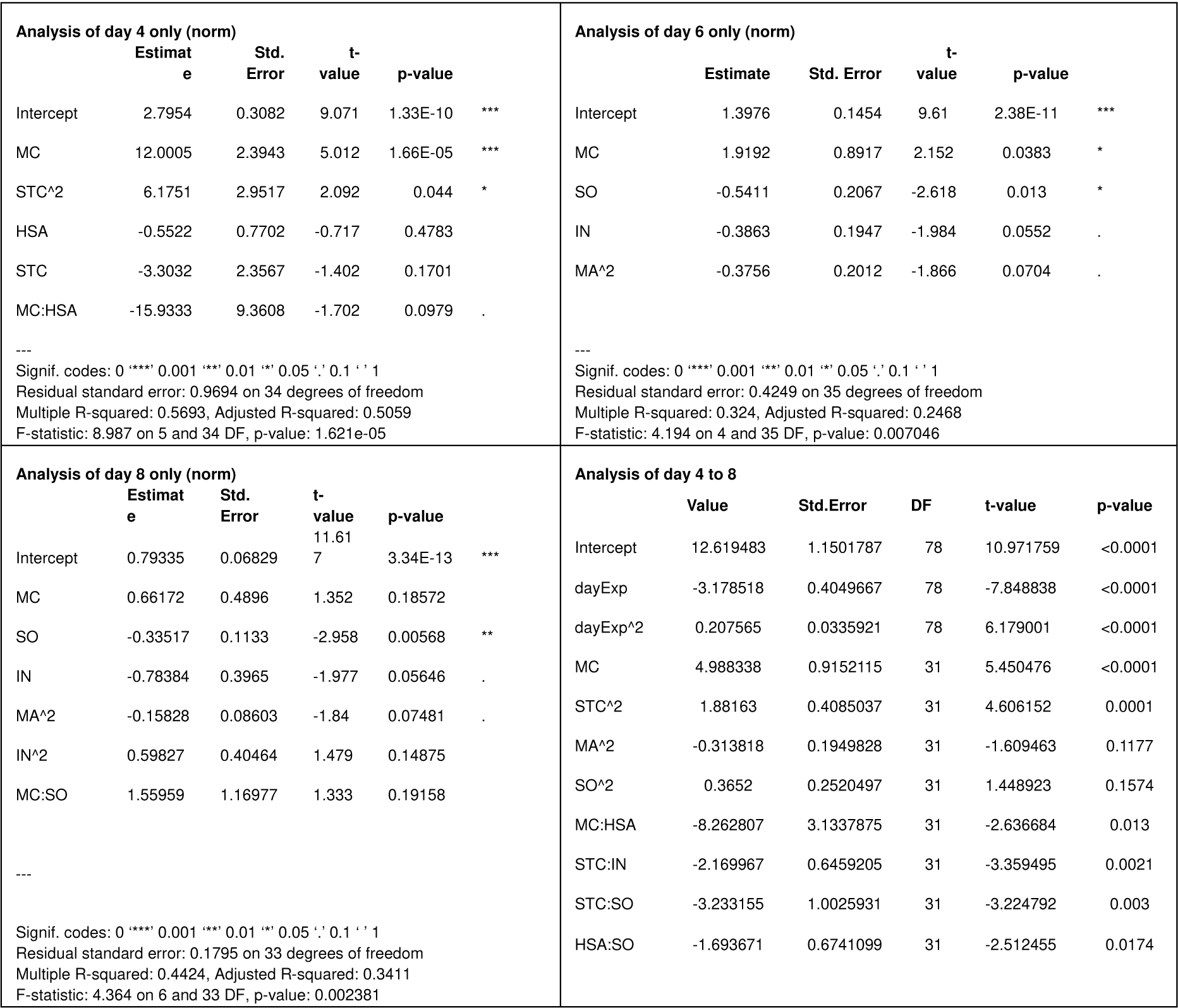
Analysis of the normalized cell culture densities.

To verify the DoE experiments, we have compared the combinations found advantageous in a standard short-term proliferation experiment (**Figure 9 A-C**). Interestingly, the combination of MC with STC in B8 has shown growth induction significantly exceeding the induction by B9 or B8+MC. This leads us to the conclusion that though DoE experiment could identify the single components, able to induce the proliferation, the identification of synergistic combinations is not possible within described setup.

**Figure 9:**
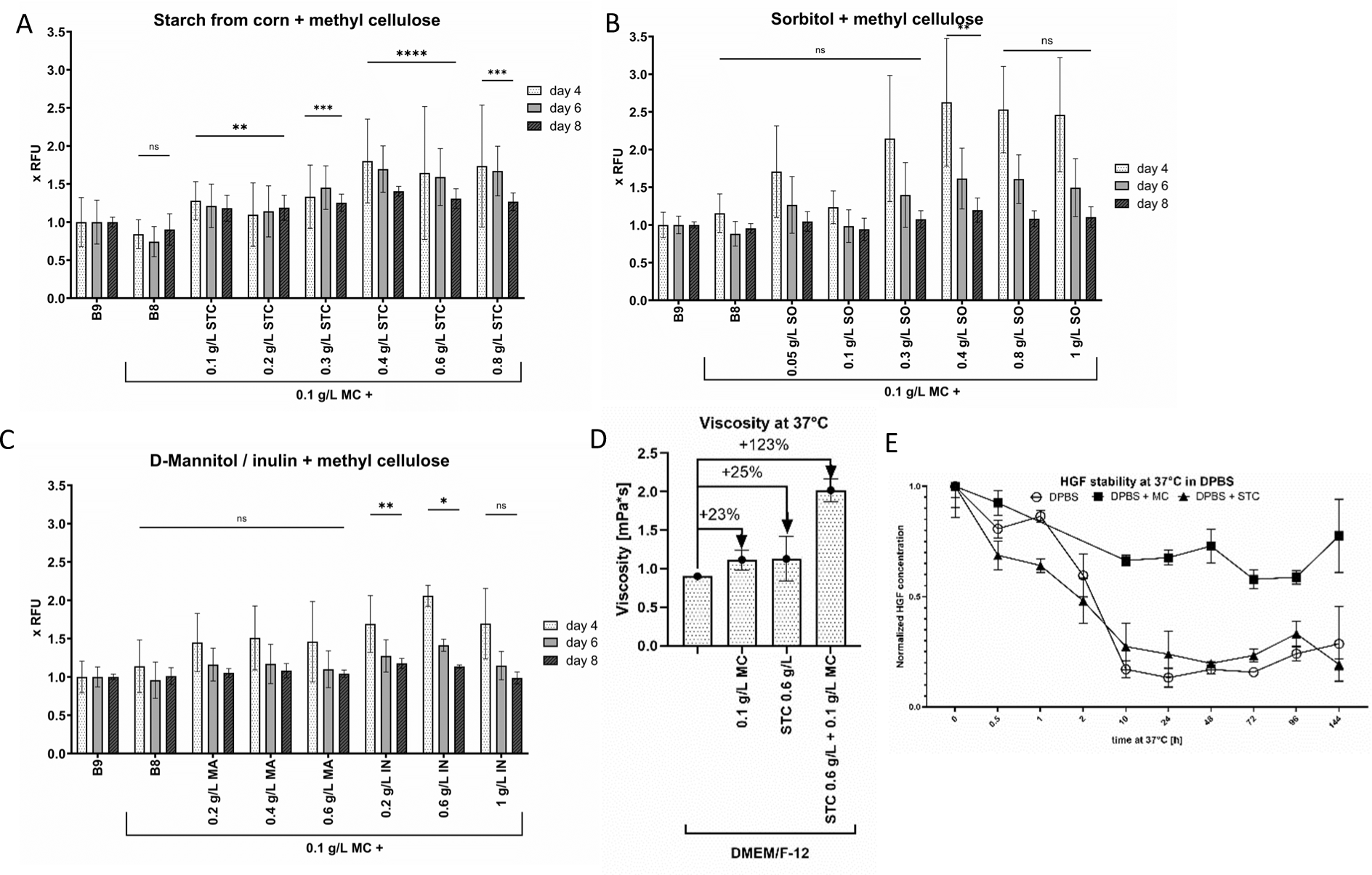
Further non-specific stabilizers with MC in B8. **(A-C)** 2000 BSCs/cm^2^ were seeded on day 0 in BSC-GM, and changed on day 1 to the designated medium. B8 with 0.1 g/L methyl cellulose (MC) and B9 medium were used as controls. Stabilizers were added to indicated end-concentrations with every medium exchange, MC always 0.1 g/L. Presto Blue assay was performed on indicated days. Obtained values were normalized to BSCs in B9. n=6 biological replicates and repeated at least twice; statistical significance was calculated by one-way ANOVA combined with Dunnett test for day 8, comparing all samples to 100% B9, and is indicated by asterisks, which are p < 0.05 (*), p <0.01 (**), p < 0.001 (***), p < 0.0001 (****). **(D)** Viscosity of the DMEM/F12 media with/without 0.1 g/L MC and/or 0.6 g/L STC was measured using Anton Paar Rheometer MCR 502 and calculated as a function of [shear stress, Pa]/[shear rate, 1/s] over 100 measuring points, each measurement except of pure DMEM/F12 was repeated trice. **(E)** Stability of rhHGF at 37°C in DPBS was measured by ELISA following manufacturer’s manuals. rhHGF in DPBS was stabilized either with 0.1 g/L MC or 0.4 g/L STC. DPBS without stabilizers was used as control. All samples were aliquoted on ice, incubated for the indicated time at 37°C, then frozen (including time point 0), and ELISA was performed after freezing the last sample.

We have further measured the viscosity of the best stabilizer mix 0.1 g/L MC + 0.6 g/L STC, which appeared approx. 100% higher than both 0.1 g/L MC and 0.6 g/L STC (**Figure 9** D), demonstrating a non-additive character of increase. This much higher viscosity could play a more significant role in higher proliferation rate of BSCs through physical properties rather than by stabilizing GFs. This conclusion was further supported by the fact that STC does not seem to demonstrate a stabilizing effect on rhHGF, in contrast to MC (**Figure 9** E).

### Long term proliferation of BSCs

One of the most important criteria to assess the efficiency of the new media are long term proliferation experiments, where cell count is monitored upon every passage and myogenic potential of the cells is assessed by calculating fusion index in differentiated cells after each passage. To validate our short-term results, we have performed such experiments for bovine satellite cells in optimized B8 and B9 media.

We conclude that additional stabilization with MC was successful in reaching higher population doublings in B8 in later passages and B9 media from the beginning, as compared to B8 and B9 without MC (**Figure 10**). BSCs’ population doublings in B9 additionally stabilized by 0.1 g/L MC, as well as doubling time were comparable with GM. Additional supplementation with rhHGF and rhPDGF did show a positive, but insignificant trend on proliferation in case of B8, but not B9.

**Figure 10:**
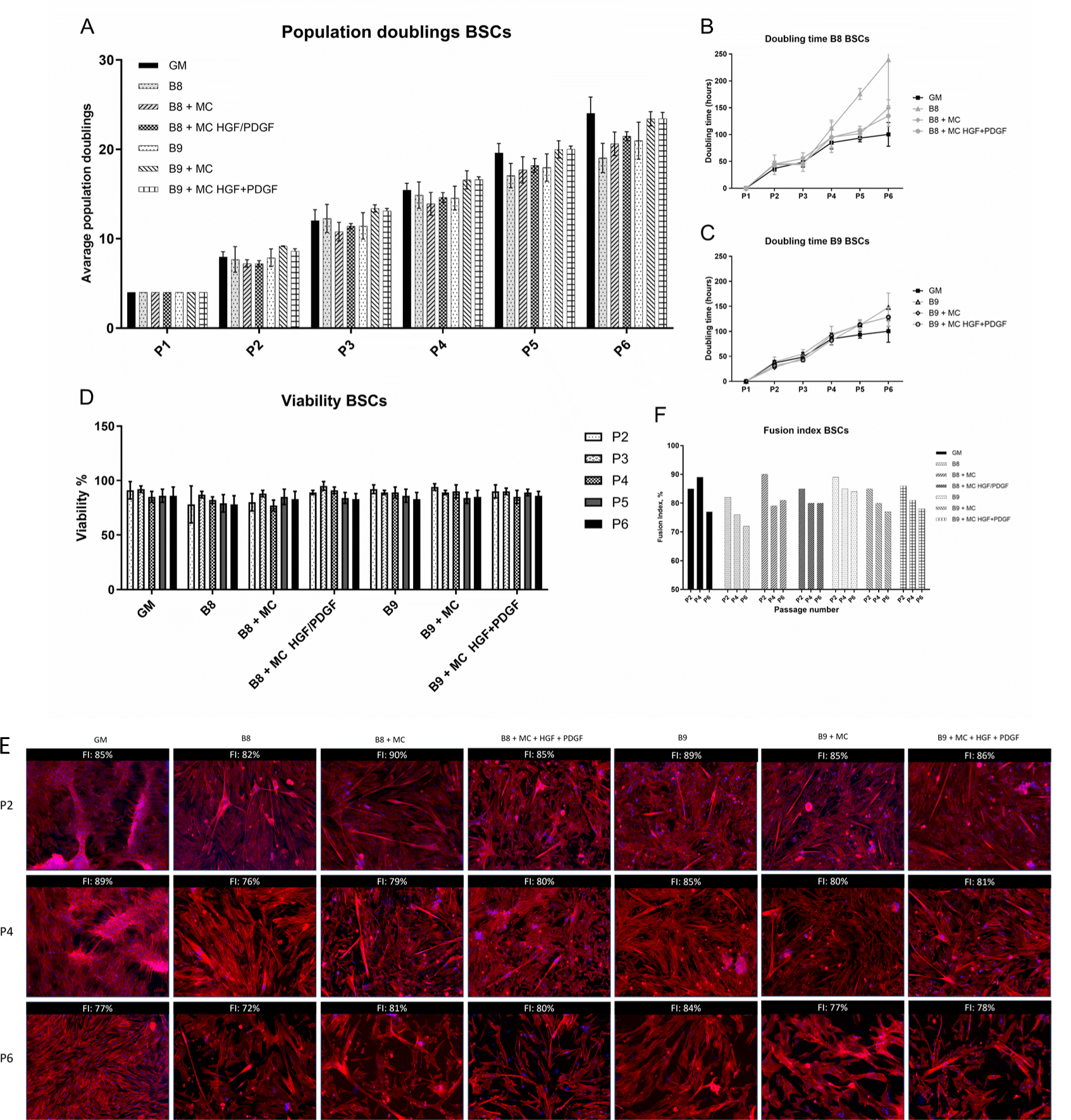
Long term proliferation experiments with BSCs in stabilized B8 and B9. Cells were cultured in 6-well plates in corresponding media, with 0.1 g/L MC and 2.5 ng/ml GFs rhHGF and PDGF added where indicated immediately upon splitting, whereas HSA was added 24h after the splitting. Cells were stained with trypan blue and counted upon passaging using Invitrogen Cell Countess, and the number of viable cells **(D)** was used to calculate population doublings **(A-C)**. Upon each passage, part of the cells was seeded for differentiation and IF staining **(E)** and fusion indexes were calculated based on the ratio of cells with at least two nuclei to all nuclei. A range of 250-1700 nuclei were counted for each sample. The count was performed by the Zeiss Zen tool for cell counting **(F)**. Immunofluorescence staining was performed for nuclei (DAPI, blue), desmin (not shown) and actin (phalloidin, red). Cells were grown to confluency and differentiated for 10 days in a serum-free differentiation medium (see Materials and Methods), with addition of stabilizers, if they were present in propagation medium. Scale bar = 100 μm. Lens: EC Plan-Neofluar 10x; Exposure Dapi: 20 ms, Phalloidin: 300 ms. GM = growth medium, FI = Fusion Index.

Best stabilizer of differentiation medium was HSA, reaching the highest fusion indexes, whereas addition of MC seemed to have a rather negative effect on differentiation efficiency of cells in HSA supplemented medium, resulting in shorter myofibers and lower fusion index. This effect is reversed in HSA-free medium, where both parameters improved upon MC supplementation (**Figure 10** E and F). These results could be explained by stabilization/retention effect MC and HSA have on different GFs from the differentiation and probably from propagation media, but would need to be further explored in more detail.

We have further investigated the potential of the other two strategies to further lower BSCs cell culture media price, which were identified in the short-term experiments: specifically, the combination of MC and STC and dilution of the growth factors to 70% from the original concentration combined with their stabilization. Addition of 0.4 g/L STC to 0.1 g/L MC could further slightly increase final cell densities in both B8 and B9 media by improving viability of the cells (**Figure 11** A-D), in B8 also promoting fusion of the myoblasts upon differentiation with no effect on fiber length or thickness, as compared to MC alone (**Figure 11** E-F). Addition of 0.1 g/L MC could as well completely compensate the dilution of the B8 media to 70% from the original protein concentrations in the long-term propagation phase, interestingly showing no additional proliferation enhancement upon 0.4 g/L STC supplementation. Also, propagation in diluted stabilized media led to thicker and longer muscle fibers upon differentiation in identical medium (e.g. B8 + MC compared to 70% B8 + MC in **Figure 11** E), leading to the assumption that GFs from propagation medium are partly stored in culture throughout medium exchange, thus decreasing the efficiency of muscle fiber maturation.

**Figure 11:**
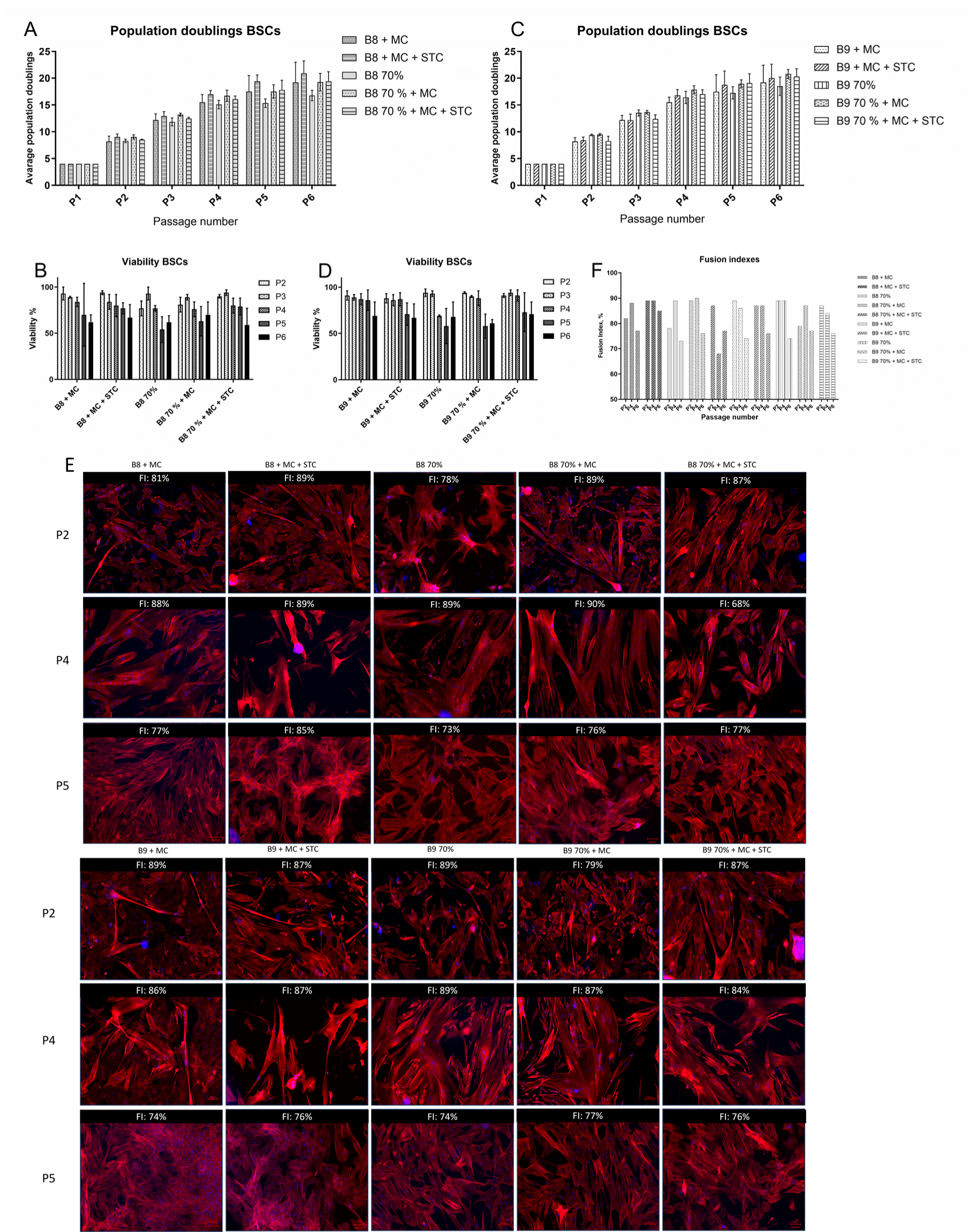
Effect of STC and GF reduction on BSCs proliferation. Cells were cultured in 6-well plates in corresponding media, with 0.1 g/L MC and 0.4 g/L STC added where indicated immediately upon splitting, whereas HSA was added 24 h after the splitting. Cells were stained with trypan blue and counted upon passaging using Invitrogen Cell Countess, and the number of viable cells **(B, D)** was used to calculate population doublings **(A, C)**. Upon each passage, part of the cells was seeded for differentiation and IF staining **(E)**, and fusion indexes were calculated based on the ratio of cells with at least two nuclei to all nuclei. A range of 250-1700 nuclei were counted for each sample. The count was performed by the Zeiss Zen tool for cell counting. **(F)** Immunofluorescence staining was performed for nuclei (DAPI, blue), desmin (not shown) and actin (phalloidin, red). Cells were grown to confluency and differentiated for 10 days in a serum-free differentiation medium (see Materials and Methods), with addition of stabilizers, if they were present in propagation medium. Scale bar = 100 μm. Lens: EC Plan-Neofluar 10x; Exposure Dapi: 20 ms, Phalloidin: 300 ms. GM = growth medium, FI = Fusion Index.

### Applicability of the stabilized medium for other relevant cell lines

Protein content of the ECM has largely formed in early chordates prior to the emergence of vertebrates^42^, leading to an assumption that the overall molecular content is also somewhat similar. We thus hypothesized that our stabilizer mixtures could probably have - to some extent - the same effect on proliferation of muscle precursor cells of other species. To investigate that, we have conducted long term proliferation experiments with porcine (PSCs) and chicken (CSCs) satellite cells in stabilized B8 and B9 in the same manner as previously described for BSCs.

Interestingly, in contrast to BSCs, the most successful proliferation medium for PSCs and CSCs appeared to be HSA free B8 + MC (**Figure 12** A-C and **Figure 13** A-C). Additional supplementation of B8 with rhHGF + rhPDGF showed a positive trend on proliferation and cell viability for PSCs (**Figure 12** A and D), but not for CSCs (**Figure 13** A-B). As in case of BSCs, HSA – supplemented differentiation medium was leading to a higher fusion index and longer muscle fibres of differentiated PSCs (**Figure 12** E-F), indicating diverging media requirements for proliferation and differentiation phases of the same cell line. Unfortunately, we were unsuccessful in growing CSCs on our imaging slides, hindering us in assessment of stabilizers’ effect on their differentiation. It should be also noted that CSCs were isolated from an adult animal and thus their media preference could significantly diverge from that of usually used chicken embryonic stem cells.

**Figure 12:**
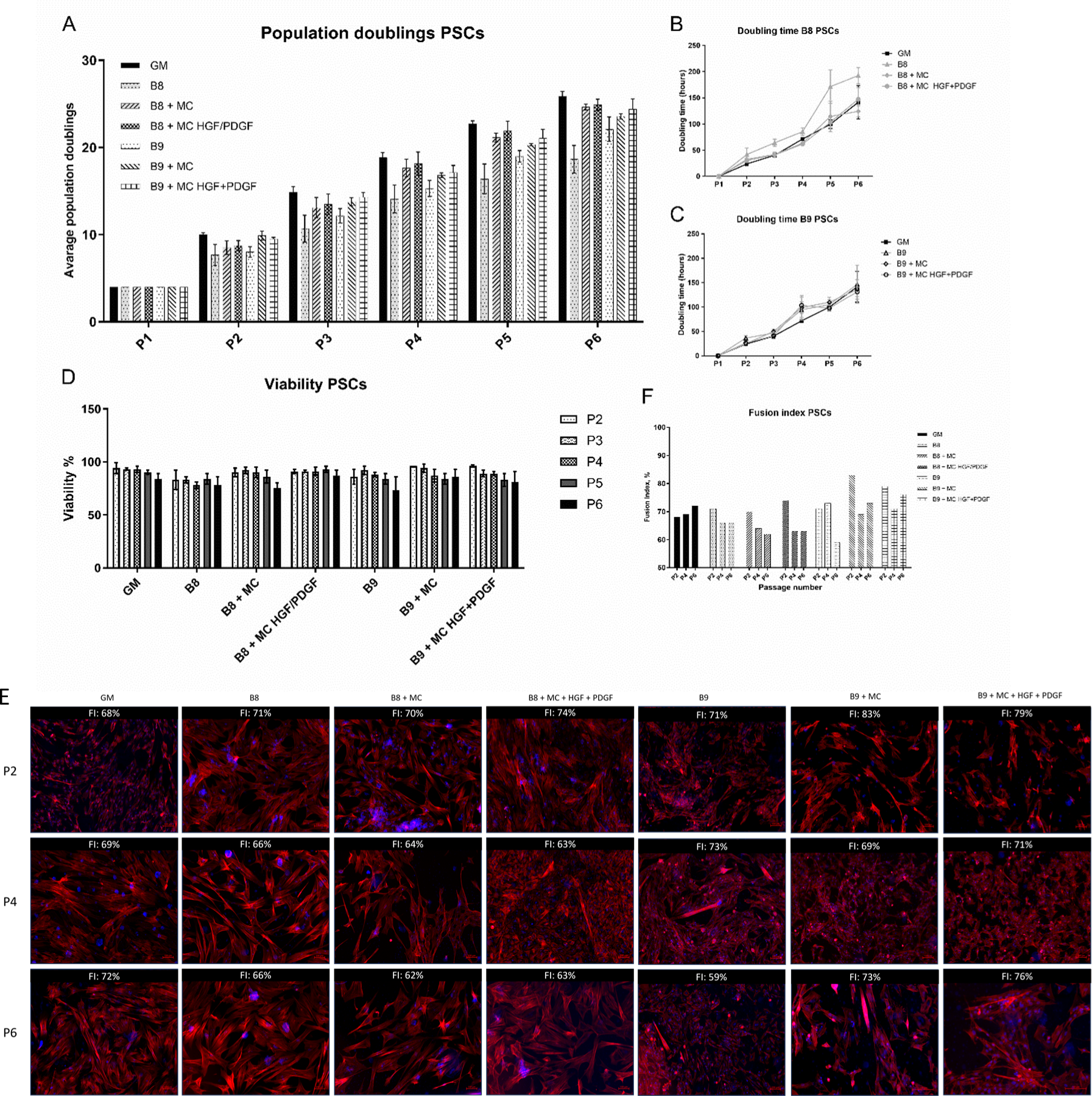
Long term proliferation experiments with porcine satellite cells (PSCs) in stabilized B8 and B9. Cells isolated from a 6 month old male *Sus scrofa domestica* were cultured in 6-well plates in corresponding media, with 0.1 g/L MC and 2.5 ng/ml GFs rhHGF and PDGF added where indicated immediately upon splitting, whereas HSA was added 24 h after the splitting. Cells were stained with trypan blue and counted upon passaging using Invitrogen Cell Countess, and the number of viable cells **(D)** was used to calculate population doublings **(A-C)**. Upon each passage, part of the cells was seeded for differentiation and IF staining **(E)** and fusion indexes were calculated based on the ratio of cells with at least two nuclei to all nuclei. A range of 250-1700 nuclei were counted for each sample. The count was performed by the Zeiss Zen tool for cell counting. **(F)** Immunofluorescence staining was performed for nuclei (DAPI, blue), desmin (not shown) and actin (phalloidin, red). Cells were grown to confluency and differentiated for 10 days in a serum-free differentiation medium (see Materials and Methods), with addition of stabilizers, if they were present in propagation medium. Scale bar = 100 μm. Lens: EC Plan-Neofluar 10x; Exposure Dapi: 20 ms, Phalloidin: 300 ms. GM = growth medium, FI = Fusion Index.

**Figure 13:**
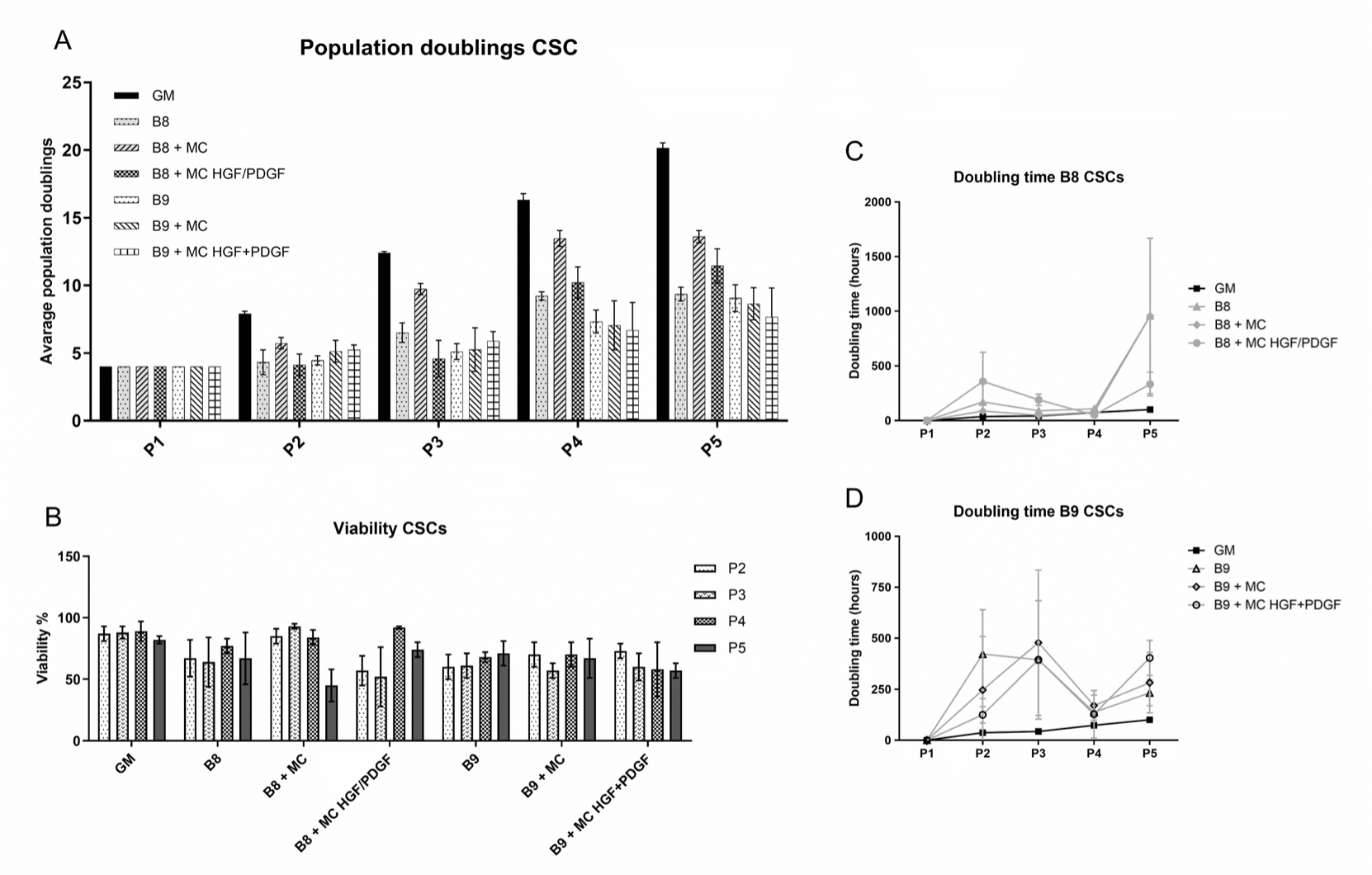
Long term proliferation experiments with chicken satellite cells (CSCs) in stabilized B8 and B9. Cells isolated from a *Gastrocnemius* of a 5 month old male rooster were cultured in 6-well plates in corresponding media, with 0.1 g/L MC and 2.5 ng/ml GFs rhHGF and PDGF added where indicated immediately upon splitting, whereas HSA was added 24 h after the splitting. Cells were stained using trypan blue and counted upon passaging using Invitrogen Cell Countess, and the number of viable cells **(B)** was used to calculate population doublings **(A)** and doubling time **(C-D)**.

Despite recent developments in serum-free culture media for mammalian cell lines, only a few studies on serum-free media exist for fish cells, majority performed in cell lines which are not relevant for cultivated fish^43,44^. Therefore, we explored the potential of stabilized B8 culture media in field-relevant cell lines such as Atlantic mackerel (*Scomber scombrus*) skeletal muscle cell line (MACK1) and European bass (*Dicentrarchus labrax*) Embryonic-like Cell Line (DLEC). As shown in supplemental figure 6, B8 medium has a higher potential to be used for MACK1 cell line, than for DLEC cell line, though we could not reach a complete substitution of FBS. For MACK1 cells, supplementation of the medium with 0.1 g/L MC and/or 0.8 g/L HSA had rather a negative effect on proliferation, though no morphological changes were detected.

### Applicability of the stabilized medium for industrially relevant cell lines

The Chinese hamster ovary (CHO) cell line established in the laboratory of Theodore Puck at the Eleanor Roosevelt Institute for Cancer Research in 1956^45^ is a well-known model cell line, widely used in genetics research and a workhorse of the biotechnological industry, famous for the production of a spectrum of recombinant products. The genetic plasticity of CHO cells, safety, scalability and compatibility with human glycosylation patterns is a testament to their popularity^46–48^. They are known as a safe production host, possessing viral entry genes, yet not expressing them^47^. Combined with compatibility with human glycosylation patterns this accentuates the qualities of CHO cells, making them eminent hosts in biopharmaceutical production, e.g. for expression of growth factors, that are highly relevant for cultivated meat area^48^.

We have therefore speculated that the ability of MC/HSA to stabilize proteins at low concentrations could also be applied in CHO cell culture to either promote cell proliferation or to stabilize the products.

An Epo-Fc producing CHO-DUXB11 line was adapted to the new media composition for 21 days. Cells were passaged to a seeding density of 0.15-0.2×10^6^ cells/mL in 10 mL working volume of supplemented growth medium every 3-4 days. Growth medium (CD CHO + 8 mM L-Gln + 0.2% ACA) was supplemented with either 0.1 g/L methyl cellulose or 0.1 g/L methyl cellulose with 0.8 g/L human serum albumin. To determine cell concentration and viability, a 500 μL sample of suspension culture was measured using trypan blue staining with the Vi-CELL™ XR cell viability analyzer (Beckman Coulter, USA). After 21 days of adaptation, cells were banked and a small-scale batch culture (20 ml total volume) from the adapted cells was performed in three biological replicates to assess whether the additives had an impact on the cell growth and viability. In addition, supernatant samples were measured using Octet RED96e (ForteBio) every 24 h to assess the titers in each condition.

Cultures growing on GM+MC showed enhanced growth and higher cell density at the end of the batch (**Figure 14 A-B**) as well as increased titers compared to the standard GM (**Figure 14** C). At the tested concentrations, none of the additives affected the binding of the EpoFc to the Octet sensors (see supplemental figure 8), which indicates that such additives would have minimal effect, if any, on the downstream processes and protein purification steps. From day 5 on (120 h) the gap in titers between the MC containing samples and GM increased, which can be explained by MC having a positive effect on the cell viability (**Figure 14** A), which resulted also in higher titers.

**Figure 14:**
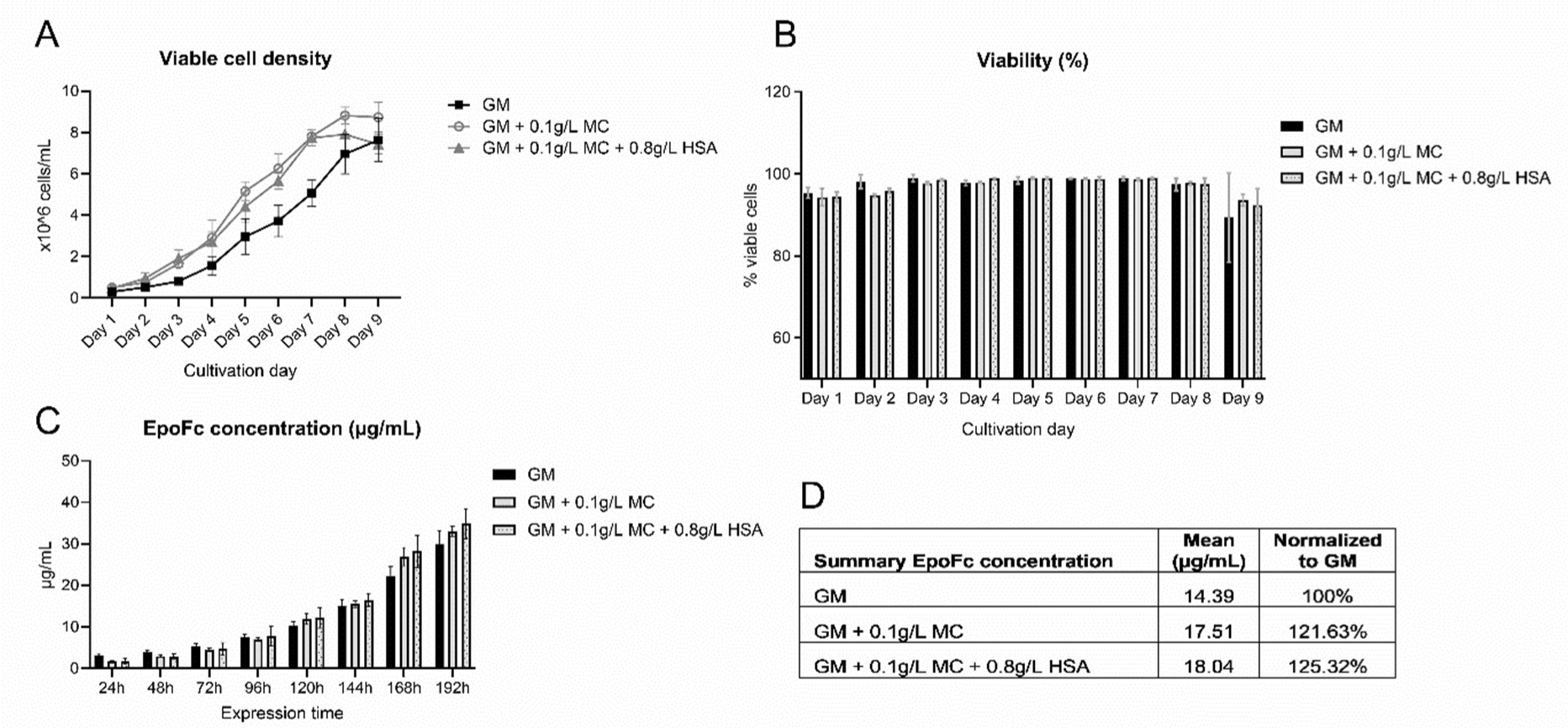
Effect of MC and HSA on expression of EPOFc in CHO-DUXB11 cell line. **(A)** Viable cell density (VCD) and viability **(B)** of CHO-DUXB11 expressing human erythropoietin fusion protein (EPOFc) in small-scale batches with the addition of stabilizers. **(C-D)** EpoFc titer concentrations from sampled supernatant of the batch cultures. Cultivations were performed in biological triplicates.

We have further applied the same strategy to Vero cells – one more industrially relevant cell line widely used as a substrate for human vaccine manufacturing^49^, derived from kidney cells of an African green monkey (AGM) started on 27 March 1962 in Chiba University in Japan. As shown in supplemental figure 6, a slightly negative effect was identified upon addition of higher concentrations of MC in B8 and commercial medium VPSFM. But, interestingly enough, B8 could successfully substitute the commercial VPSFM medium, which is substantially more expensive, and the dilution strategy have demonstrated that both 80% and 70% of VPSFM promoted significantly higher proliferation rates, than 100% VPSFM.

## Discussion

Development of the serum free medium for expansion of primary satellite cells has come to a new level in the last two years, as several medium compositions were presented demonstrating efficiency comparable to FBS containing medium compositions. This was facilitated by observing the importance of using not only the right combination of GFs, but also after introduction of medium stabilization, mimicking to a certain degree the ECM content. Stout *et al.* ^50^ and Kolkmann *et al*.^6^ use well known and very widely applied human or bovine serum albumin for medium stabilization, Skrivergaard *et al.*^9^ also, but in combination with Fetuin. In case of B9 medium containing 40 ng/mL FGF-2, about 61% of the whole B9 medium costs are attributed to HSA (see supplemental table 2, basic medium – DMEM-F12 – not considered). In the medium formulation of Kolkmann *et al.,* 5 g/L HSA are used, which, in combination with another ECM protein fibronectin at 10 μg/mL, sums up to approx. 475 USD/L, or 77% of the costs only for stabilization^6^. And, finally, Skrivergaard *et al.* make use of a combination of 600 μg/mL Fetuin (fetal variant of HSA) and 75 μg/mL BSA, resulting in staggering 81% of the media costs attributed to EMC mimetics – due to a high price on pharmaceutical grade Fetuin^9^.

ECM mimetics contribute to the vast majority of the medium price for the three mentioned media compositions (**Figure 15**), posting a new challenge for cultivated meat industry, as they are not only pricy, but also applied in high concentrations (see supplemental table 2 for comparison). Recent evaluations of recombinant protein costs and volumes necessary for cost-competitive cultivated meat production suggest that 96.6% of production volume is expected to be attributable to albumins, and only under 4% will comprise transferrin, insulin, growth factors and other proteins^10^. Based on these assumptions, to produce cultivated meat in volume of 1% of the global meat market would require millions of kilograms of albumin, which by far exceeds the current production volumes of many recombinant industrial enzymes, and of the serum derived albumins as well. In a recent paper Stout *et al*. presented the possibility to substitute HSA with seed protein isolates, which are not yet commercially available, but can be produced in-house^30^. The costs of this solution are yet to be evaluated, including the requirement of storage at -80°C for the isolates to stay biologically active.

**Figure 15:**
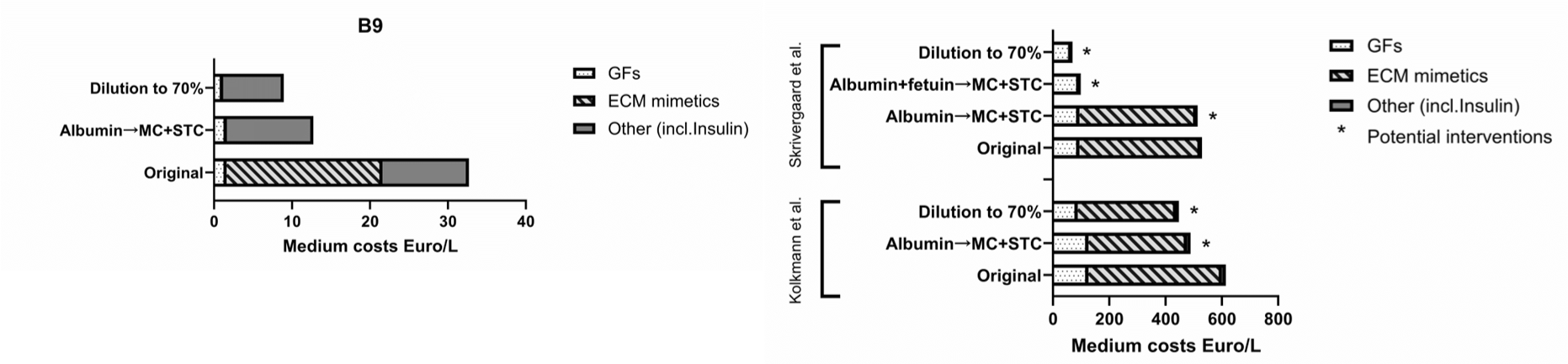
The potential of alternative protein stabilizers and of dilution strategy to reduce medium costs. Medium cost calculations are based on bulk prices summed up in supplemental table 2. *Potential interventions – estimation of the potential of media cost reducing strategies identified in this study, not evaluated in laboratory conditions.

But although albumins are very widely used for non-specific stabilization of biologically active proteins^19–21^, they are far from being the only known protein stabilizers – salts, sugars, amino acids and hydrogels are also used for this purpose, some of them known under the term “chemical chaperones”, and are usually utilized for refolding of recombinant proteins from inclusion bodies, or for storing proteins over longer time periods^12,22,51^. The greatest limitation of stabilization approach is its’ specificity - it is recognized that various stabilizers may exert differing degrees of influence on different growth factors; also, accurately predicting these effects remains an unresolved challenge^21,24^. But, in our opinion, such ability of some low-cost chemicals to drastically enhance the stability of the most expensive cell culture medium components has a great potential in cultivated meat area, and probably also in industrial production of biopharmaceutics.

We have therefore chosen several stabilizers, which are readily available in big quantities, are low-priced in production and storage, and possess extremely long shelf-lives. We have narrowed down to several components based on the published literature, focusing on stabilization of the GFs present in B8/B9 media, and complemented them with potential stabilizers based on structural similarity to stabilizers known to effectively prolong half-life of crucial GFs (**Table 2**). We have tested these stabilizers and their combinations in low concentrations in short- and long-term cultivations on BSCs, and assessed their influence on differentiation. We have further tested the carry-over of the identified conditions onto other cultivated meat/fish relevant cell lines (PSCs, CSCs, DLEC, MACK1) as well as onto industrially relevant cell lines (CHO, Vero) and their effect on cell line relevant characteristics.

In this study we have shown that a row of known and new protein stabilizers and their combinations can be used in cell culture – methyl cellulose (MC), DL-alanine (ALA), starch from corn (STC), etc. ^24,52^ – instead of HSA for the comparable stabilization effect, and that the final combination of stabilizers is cell line specific. Our results are in line with the FGF-2 stabilization study from Benington *et al*., who demonstrated that a combination of two stabilizers – MC + ALA or MC + HSA – performs much better than any component alone^24^. In our case, superior effect of the combinations could be based on the macromolecular crowding setup we had in our experiments, where high concentrations of largely “inert” macromolecules (MC + HSA or MC + STC) are mimicking the ECM environment, with its limited space containing only considerably restricted amounts of free water^53^. It is also known that MC, as well as other polymeric stabilizers such as PEG, sequesters hydrophilic proteins, preventing their aggregation and subsequent decay, and in some of our setups proteins are additionally stabilized by HSA through ionic, electrostatic and hydrophobic interactions^19,21^. We hypothesize that MC, structurally resembling glycosaminoglycan chains of ECM proteoglycans^29^, together with high concentration of HSA, could reasonably mimic the ECM – better than MC or HSA alone. We also applied DL-ALA in the present study as amino acids with no net charge, e.g. alanine and glycine, are often used as cryoprotectants of proteins, providing stability through weak electrostatic interactions^54^.

The best hits among known and potential stabilizers both in short- and long-term cultivations were methyl cellulose (MC) and starch from corn (STC). High concentrations of MC (registered as E461 in the EU) are traditionally used in various cell culture methods, such as preservation of cell function in suspension^55^, cell aggregation for 3D spheroid formation^56^, or colony forming cell assay^57^, where concentrations from 3 to 0.2% methyl cellulose are used. It is also widely used in food industry as thickener, emulsifier and for its binding and gelling properties – e.g. in plant-based meat alternatives^37^, that was recently criticized due to its synthetic nature^37^, as it is manufactured by heating cellulose with a caustic solution (such as sodium hydroxide) and subsequently treating it with methyl chloride^38^. STC, on the other hand, is a naturally occurring edible polymer with the same various applications in the food industry as thickener, stabilizer and binder, is also used as a hydrogel-polymer^58^ and a drug delivery system^59^. Although we could show that both of the substances were exerting a significant positive effect on BSCs proliferation, and in case of MC the stability of HGF was greatly improved, there seemed to be no specific binding of the GFs and stabilizers, as it is known for other well described stabilizers of GFs, e.g. heparin^12^. We thus conclude that the MC mechanism of action, as well as that of STC, might be based on the crowder’s effect, leading to a longer half-life of some proteins, as it is known e.g. for the favorite molecule of the nanomedicine and another broadly used chemically manufactured food and cosmetics additive – polyethylene glycol (PEG)^60^, but could additionally be a result of the improvement of cell adhesion through increase in media viscosity.

Interestingly, for both porcine and bovine satellite cells, B8 was the worst performing, but still a possible long term cultivation medium, whereas Stout *et al*. have reported, that it was impossible to cultivate their bovine satellite cells for longer than two passages^5^. It might be due to the sourcing discrepancy between the cells – we were unable to source samples from 1 to 2 month bulls, and were constrained to using cells from approx. 1.5 years old bulls instead. This might have led to a higher flexibility regarding media requirements of the more mature satellite cells, but can have other disadvantages – for example, higher doubling times in the later passages, than those reported for the satellite cells from 1 to 2 month bulls – for the P5 cells doubling times of approx. 75 h if sourced from an older animal, as compare to around 50 h, if sourced from a younger animal^5^. This is in line with other reports on the significantly higher doubling time of the satellite cells’ isolated from older animals – e.g. mice^61^, pigs^62^ and humans^63^.

The main goal of this work was to establish strategies for lowering the price for cultivation medium. Stabilizing the media with newly identified ECM-mimicking components allowed us not only to substitute HSA, but also to reduce the initial concentration of B9 components to 70% from the original without losses in BSCs cell density, leading to a 73% combined price reduction and a more efficient muscle fiber maturation. Our results underscore the importance of the ECM-mimicking protein stabilizing compounds and this line of research should be continued to substitute or lower the concentration of further expensive media components, such as fetuin and fibronectin – both comprising a majority of the recently published serum-free fully chemically defined medium formulations (**Figure 15**). Still, it is to be validated that approx. 20% higher viscosity, caused by the addition of single stabilizers, does not lead to a substantial raise in energy consumption in stirred bioreactors, and does not affect heat distribution, leading to a less efficient cooling. Also, prediction of the stabilizers’ effect on a specific GF remains out of reach, and stabilizers influence on different growth factors is quite specific, complicating transmission of the media optimization results onto other cell lines^21,24^. This was also confirmed in our study, as most efficient stabilizer combinations for BSCs, PSCs and CSCs diverged from each other, and none of the final tested combinations was successful in improving fish cell lines’ proliferation, or proliferation of Vero cell line.

Other very costly components of the media could be the coating proteins (such as iMatrix, bovine collagen type I or vitronectin, as used in this study), which are usually necessary for optimal adhesion of primary satellite cells in serum-free media. This cost factor was not considered in this work, as the employment of suspension-suitable cell lines or cultivation in cell aggregates upon upscaling seems to be the cultivation of choice for most of the companies due to higher efficiency of medium utilization, which makes coating proteins unnecessary.

To sum up, this and further research will eliminate the bottleneck of a high albumin dependency of the cultivated meat industry and will allow a faster upscaling of the technology without a need to make massive investments into additional infrastructure for recombinant albumin production. Our findings can also be transferred to medical research to support the efforts of 3R concept of minimize animal use (Replacement, Reduction, and Refinement)^64^ by producing a more cost-effective media and raising the reproducibility of cell culture experiments. Another line of possible application is in pharmaceutical industry in production of various therapeutical proteins, as shown in this study on the example of EpoFc production in CHO-DUXB11 cell line.

## Materials and methods

### Materials

rhHGF ELISA kit: Thermo Fisher #BMS2069INST; rhPDGF ELISA kit: Thermo Fisher #BMS2071. Monolith His-Tag Labeling Kit RED-tris-NTA 2nd Generation kit (Cat#MO-L018). B8 (HiDef-B8 500X) was purchased from Defined Bioscience (#LSS-201). For sourcing GFs, hormones and cytokines see supplemental table 1, for the sourcing of stabilizers - **Table 2**.

### Isolation of satellite cells

Bovine Satellite Cells (BSC) were isolated from a 19 months old, castrated Simmental Ox (*Bos taurus*) *M. semitendinosus* muscle tissue sample provided by Marcher Fleischwerke GmbH. Porcine Satellite Cells (PSCs) isolated from a 6 month old male *Sus scrofa domestica M. semitendinosus* muscle tissue sample provided by Marcher Fleischwerke GmbH. CSCs = Chicken Satellite Cells isolated from a *Gastrocnemius* of a 5 month old male rooster provided by Thomas Raberl. Cells were isolated from sacrificed animals using a standard protocol described e.g. by Stout *et al*.^65^

A sample the size of ∼0.5 g of skeletal muscle tissue was extracted. The muscle probe was transported in transport medium on ice and immediately further processed, where it was cut up into small pieces and digested in 0.2% collagenase II (Worthington Biochemical #LS004176, Lakewood, NJ, USA; 275 U/mg) for 45-60 min until the paste was homogenized. Digestion reaction was brought to a halt with BSC growth medium (P-GM). The cells were filtered twice, counted using an Invitrogen Countess Automated Cell Counter and plated at a density of 100,000 cells/cm^2^ onto uncoated tissue-culture flasks. Throughout the incubation over 24 h at 37 °C with 5% CO_2_ adherent cells attached at the surface of the tissue-culture flask and SCs stayed in suspension and were transferred to coated flasks at a density of 2000 cells/cm^2^ with 1.5 µg/cm^2^ recombinant human Vitronectin (FisherScientific #15134499). SCs were left untouched for three to four days before P-GM was changed every two days until a maximum of 70% confluence was reached and the cells were frozen in FBS with 10% dimethyl sulfoxide (DMSO, Sigma #D2650) or passaged for screening or differentiation by using 0.25% trypsin-EDTA (ThermoFisher #25200056).

### Confirmation of SCs identity and differentiation

After isolation, undifferentiated SCs were identified by staining for Paired-box 7 (Pax7) marker. SCs were cultured in BSC growth medium (BSC-GM) on a cover glass until they reached 70% confluence, fixed with 4% paraformaldehyde (FisherScientific #AAJ61899AK) for 30 min, washed in DPBS, permeabilized for 15 min with 0.5% Triton-X (Sigma #T8787) in DPBS, blocked for 45 min with 5% goat serum (ThermoFisher #16210064) in DPBS with 0.05% sodium azide (Sigma #S2002), and washed with DPBS with 0.1% Tween-20 (Sigma #P1379). Primary Pax7 antibodies (ThermoFisher #PA5-68506) were added at a dilution of 1:500 in blocking solution (Aligent Antibody Diluent #S202230-2) containing 1:100 Phalloidin 594 (ThermoFisher #A12381) to cells and incubated at 4 °C overnight. Further, cells were washed with DPBS + Tween-20 and incubated with secondary antibodies for Pax7 (ThermoFisher #A-11008, 1:500) for 1 h at room temperature, washed with DPBS + Tween-20 and mounted with Fluoroshield mounting medium with DAPI (Abcam #ab104139). Visualization and imaging were performed by the Core Facility Imaging at Medical University Graz to validate the satellite cell purity of the isolated cell population.

Isolated SCs were differentiated for ten days. After growing cells to confluency in BSC-GM, the medium was changed to Differentiation Medium (DM) and then 50% of the medium was regularly changed (every two to three days) until SCs reached differentiated state and were fixed and prepared as previously described. Phalloidin 594 was diluted 1:100 in blocking solution (Aligent Antibody Diluent #S202230-2) and incubated at 4 °C overnight. Fluoroshield mounting medium with DAPI was mounted next day.

### Media composition

Skeletal muscle tissue sample was transferred into transport medium consisting of DMEM + Glutamax (ThermoFisher #10566016) and 1% Antibiotic-Antimycotic (ThermoFisher #15240062) after extraction. For Isolation of BSCs, Primocin growth medium (P-GM) was used containing DMEM + Glutamax (ThermoFisher #10566016), 1% Primocin (Invivogen #ant-pm- 1), 20% fetal bovine serum (FBS; ThermoFisher #26140079) and 1 ng/mL human FGF basic/FGF2/bFGF (R&D Systems #233-FB-025/CF). After passage one, P-GM was changed to growth medium (GM) where Primocin is switched out with 1% Antibiotic-Antimycotic (ThermoFisher #15240062). Serum free differentiation medium was prepared according to Messmer *et al*.^66^

For short-term and long term growth analysis, cells were plated in 96-well (Greiner #655180) or 24-well tissue culture plates (Greiner #662160) containing BSC-GM with 1.5 µg/cm^2^ recombinant human Vitronectin (FisherScientific #15134499). After incubation for 24 h, cells were washed with DPBS (ThermoFisher #14190250) and BSC-GM was changed to screening medium - B8 medium containing DMEM/F12 (ThermoFisher #11320033), HiDef-B8 medium aliquots (Defined Bioscience #LSS-201) and 1% Antibiotic-Antimycotic, supplemented or not supplemented with 0.8 g/L HSA. According to the screening method, defined concentrations of medium components (Suppl. Table 1) and stabilizers (Table 3) were added. All added medium components were reconstituted and diluted in DPBS containing 0.8 g/L HSA (Sigma #A9731-1G), highly concentrated aliquots were stored at -80°C.

### Screening of BSCs/PSCs/CSCs

BSCs/PSCs/CSCs were thawed in 5 mL GM and centrifuged at 200 x g for 3 min. The supernatant was discarded, and the cells were washed with 5 mL DPBS. After another centrifugation step and removal of the supernatant, the cells were carefully resuspended in 5 ml BSC-GM and counted with Invitrogen Countess Automated Cell Counter.

For short term analysis SCs (800 cells/well = approx. 2000 cells/cm^2^) were plated in 96-well tissue culture plates (Greiner #655180) containing 100 µl BSC-GM with 1.5 µg recombinant human Vitronectin (FisherScientific #15134499)/cm^2^. After 24 h of incubation (day 1) at 37°C and 5 % CO_2_, the cells were rinsed with DPBS (ThermoFisher #14190250), BSC-GM was removed, screening medium (B8 or B9) and screening components were added (see the list of components and their sourcing in supplemental table 1). On indicated days Presto Blue assay was performed. On day 3 after Presto Blue assay screening components were added again into the fresh medium. Over the weekend cells were covered with 200 µl medium, instead of 100 µl. On the last day, the cells were rinsed with DPBS and frozen at -80°C.

Long term analysis was performed in 24-well plates (Greiner #662160) containing 1 mL BSC-GM with 1.5 µg recombinant human Vitronectin (FisherScientific #15134499)/cm^2^. After 24 h of incubation (day 1) at 37°C and 5 % CO_2_, the cells were rinsed with DPBS (ThermoFisher #14190250), BSC-GM was removed and screening medium with or without screening components/stabilizers were added. At 60-70% confluency cell were counted using Invitrogen Cell Countess and the total viable cells of three replicates were further analysed. Cells were then passaged using Trypsin-EDTA (0.25%) (ThermoFisher #25200056) and plated in either growth medium or B8 along with 1.5 µg recombinant human Vitronectin /cm^2^. After 24 h stabilizers and screening components were added, and cells were analyzed for proliferation and differentiation.

### Cultivation of CHO-K1 cell line

The CHO-K1 cell line was obtained from the ECACC collection (UK) and the CHO-DUXB11 EpoFc from Taschwer *et al*.^67^ Cells were kept in suspension culture in an orbital shaking CO_2_ incubator at 7% CO_2_, 37°C and 220 rpm in TPP^®^ TubeSpin bioreactor tubes (Merck, USA). Cells were passaged to a seeding density of 0,15-0,2×10^6^ cells/mL in 10 mL working volume every 3-4 days. To determine cell concentration and viability, a 500 µL sample of suspension culture was measured using trypan blue staining with the Vi-CELL™ XR cell viability analyzer (Beckman Coulter, USA).

For the CHO-K1 cell line CD-CHO medium (Gibco™, Carlsbad, CA, USA) supplemented with 0.2% Anti-clumping agent (Gibco™, Fisher Scientific, USA) and 8 mM L-Glutamine (Sigma Aldrich, USA) was used as the standard Growth media (GM). For the CHO-DUXB11 EpoFCcell line CD-CHO medium (Gibco™, Fisher Scientific, USA) supplemented with 0.2% Anti-clumping agent (Gibco™, Fisher Scientific, USA), was used as the standard GM.

### Presto Blue assay

To measure the metabolic activity, which correlates with the number of cells, Presto Blue assay was performed in 96-well plate format according to the instruction manual. Shortly, 10 µL Presto Blue reagent (ThermoFisher #A13262) was added to the remaining 90 µl of medium in each well and incubated at 37°C for 1 h. After incubation, the medium was transferred to a fresh 96-well plate (Greiner #655101) and the fluorescence at 560 nm (excitation) and 590 nm (emission) was measured with a plate reader (BioTek SynergyMx). Depending on the time point of the assay, cells were covered with 100 or 200 µL of fresh B8 or B9 medium and further incubated.

### Hoechst assay

In order to verify Presto Blue assay data, we used Hoechst 33258 assay to assess the quantity of DNA per well, which directly correlates with the cell number.

On the last day of every screening, the screening plates were frozen at -80°C (see Screening). After thawing to room temperature, 100 µL ddH_2_O were added per well and the plates were incubated at 37°C for 1 h. The plates were then again frozen at -80°C and thawed to room temperature. 100 µL of Hoechst dye reagent - consisting of 25 µL Hoechst 33258 stock solution (10 mg Hoechst 33258 (ThermoFisher #H1398)/mL in DMSO + ddH_2_O (1:4 v/v)) in 10 mL TNE buffer (10 mM Tris, 2 M NaCl, 1 mM EDTA, 2 mM sodium azide, pH 7.4, were added. After 15 min of incubation at room temperature, the fluorescence was measured at 352 nm (excitation) and 461 nm (emission).

### NanoTemper Thermophoresis assay

Monolith His-Tag Labeling Kit RED-tris-NTA 2nd Generation kit (Cat#MO-L018) was used to label HIS-tagged rhHGF via binding to its His-tag according to the instruction manual. Thermophoresis assay was performed at 25°C with 25 nM HIS-tagged rhHGF with MC in a range of concentrations from 0.25 g/L to 0.00032 mg/L.

### Statistical analysis

For statistical analysis of Presto Blue and Hoechst assays, the data was tested for significance using ordinary one-way ANOVA followed by Dunnett’s multiple comparisons test was performed using GraphPad Prism version 10.2.3. Statistical significance was reached when p < 0.05 (*), p < 0.01 (**), p < 0.001 (***) and p < 0.0001 (****). Non-significant results (p ≥ 0.05) were marked as “n.s.”. The error bars in the graphs represent the standard deviation.

### DoE

The design space is spanned by the 6 stabilizers given in **Table 3**. Trials of the experiment were only able to be run at the grid points within the design space specified by the 8 stabilizer concentrations (see **Table 3**). A space-filling design^39^ with 48 trials was created and each trial was shifted to the nearest grid point afterwards. The resulting concentrations of each of the 6 stabilizers were summed up and the 8 trials with the largest sums were excluded from the design to avoid possible advert effects of higher viscosity.

### Linear and linear mixed effects models

For each day (4, 6, and 8) separately we fitted linear second order regression models with the 6 stabilizers as explanatory variables. The largest models could contain all 6 stabilizers, their quadratic terms and their two-way interactions as regressors. The best model was chosen by a stepwise forward regression using Akaike’s information criterion (AIC) to decide whether an additional regressor enters the model or to stop the estimation.

To analyze the measurements of days 4-8 together in a single model we fitted a second order linear mixed effects model to our data additionally including the day and its quadratic term as regressors. Again, the best model was chosen by a stepwise forward regression using AIC.

## Author contributions

Lisa Schenzle: investigation, visualization, data analysis, draft writing. Kristina Egger: investigation, visualization, data analysis, draft writing. Bernhard Spangl: DoE planning and data analysis. Mohamed Hussein and Nicole Borth: investigation, data analysis. Atefeh Ebrahimian and Harald Kuehnel: investigation, data analysis. Frederico C. Ferreira and Diana M. C. Marques: investigation, data analysis. Beate Berchtold: investigation, visualization, data analysis. Aleksandra Fuchs: conceptualization, resources, writing-reviewing and editing, supervision, funding acquisition. Harald Pichler: resources, reviewing and editing, supervision.

## Acknowledgements

We are very grateful to DFK foundation, Torggler Rechtsanwälte GmbH and their scientific advisory board Dr. Kurt Schmidinger and Prof. Tomislav Cernava for the opportunity to work on this exciting project and for their valuable input. The project was performed at COMET center acib: Next Generation Bioproduction, and is additionally funded by BMK, BMAW, SFG, Standortagentur Tirol, Government of Lower Austria and Vienna Business Agency in the framework of COMET - Competence Centers for Excellent Technologies. The COMET Funding Program is managed by the Austrian Research Promotion Agency FFG. We very heartily thank Andrew Stout for his support of this work and for his advice, Jörg Mai from Marcher Fleischwerke GmbH and Thomas Raberl for providing access to animal tissues. We are very thankful to Esther Föderl-Höbenreich and Daniela Pabst from Graz Medical University for the help in establishing satellite cell isolation protocol, and to Zerina Arnautalic for her dedication and support in conducting the lab-work for the production and analysis of the EPO-FC fusion protein described in this work. We also thank Good Food Institute for their valuable comments and suggestions. Additional appreciation is directed to VIZE & Tom Gregory for inspiration.

