## Supplemental data for "Never let me down: new possibilities for lowering serum free cultivation media costs"

### Title

### *Co-first authors

### #Correspondence

**Supplemental table 1:** Overview of the components screened for induction of higher proliferation rates in primary BSCs.

| Item | Response to | Inflammation | Article Nr | Range published / used in our screening | Positive effect published |
|---|---|---|---|---|---|
| **Growth factors / Injury signals** | | | | | |
| rh IGF1 | injury | developmental | ab9573 | 0-250 ng/mL | Yoshida 2020[1] |
| rh HGF | injury | anti-inflammatory | ab245957 | 0-250 ng/mL | Anderson[2] |
| rh IL-1α | injury | pro-inflammatory | PeproTech #200-01A | 0-250 ng/mL | Fu 2015[3] |
| rh IFN-γ | injury | pro-inflammatory | PeproTech #300-02 | 0-250 ng/mL | Fu 2015[3] |
| rh TNF-α | injury | pro-inflammatory | PeproTech #300-01A | 0-250 ng/mL | Fu 2015[3] |
| rhPDGF-BB | injury | | Genscript #Z02529 | 0-250 ng/mL | Wang 2019[4] |
| **Hippo pathway inhibitors** | | | | | |
| Lysophosphatidic acid | Hippo | Pro-inflammatory | Sigma L7260-1MG | 10 or 25 µM | Memphis Meat Patent[5] |
| Estrogen (17β-Estradiol) | Hippo | anti-inflammatory | Sigma E2758-250MG | 3 ng/mL | Mangan 2014[6]<br>Kamanga-Sollo 2014[7] |
| Dihydrotestosterone | Hippo | anti-inflammatory | Sigma NMID680 | 3 ng/mL | Johnson 1996[8] |
| rm Wnt3a | Hippo | | 1324-WN-002 | 250 ng/mL | Park 2015[9] |
| rh Wnt-5b | Hippo | | R&D 7347-WN-025/CF | 125-750 ng/mL | Park 2015[9] |
| **Myokines** | | | | | |
| rh YKL-40/CHI3L1 | sport | anti-inflammatory | ab182706 | 0-250 ng/mL | Görgens 2016[10] |
| rh GASP-1 | injury + sport | | Sigma SRP6220-10UG | 2.5-250 ng/mL | Brun 2014[11] |
| rh Follistatin | injury + sport | anti-inflammatory | ab50163 | 6–600 ng/mL | Gilson 2009[12] |
| rh IL-6 | injury + sport | pro-inflammatory | Biorad PBP021 | 0.01 - 250 ng/mL | Stout 2021[13] |
| rh IL-4 | injury + sport | anti-inflammatory | Peprotech #200-04 | 2.5-250 ng/mL | Chang 2019[14] |
| rh IL-13 | injury + sport | anti-inflammatory | Peprotech #200-13 | 0-250 ng/mL | Fu 2015[3] |
| rh IL-15 | sport | | Peprotech #200-15 | 0-250 ng/mL | Lightfoot 2016[15] |
| rh LIF | sport | developmental | Peprotech #200-16 | 0-250 ng/mL | Maleiner 2018[16] |

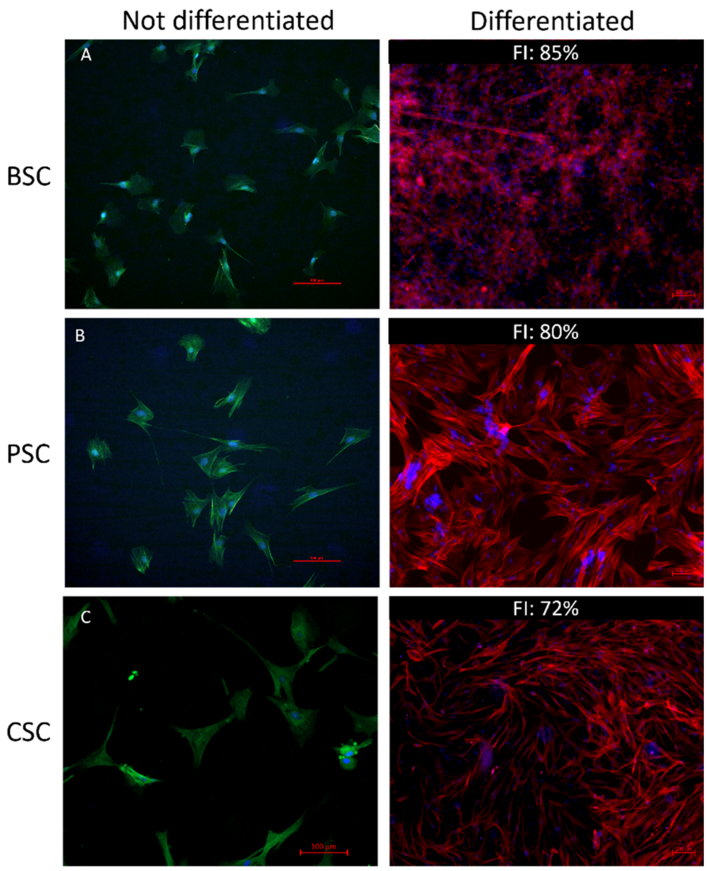

**Supplemental figure 1:** Differentiated and non-differentiated cell from different species after isolation. BSC = Bovine Satellite Cells; PSC = Porcine Satellite Cells Cells isolated from a 6 month old male *Sus scrofa domestica*; CSC = Chicken Satellite Cells isolated from a Gastrocnemius of a 5 month old male rooster. Cells were isolated from sacrificed animals using a standard protocol described e.g. by Stout et al. [17] Satellite cell (SCs) were propagated for several days after isolation, and stained for the typical marker of non-differentiated satellite cells PAX7 labelled with Alexa Fluor™ 488 (green); or differentiated as described in the methods section and stained for the typical marker of differentiated satellite cells - actin using phalloidin labelled with Alexa Fluor™ 594 (red). Nuclei were visualized using DAPI (blue). FI – fusion index calculated as a percentage of nuclei within the multinucleated myotubes, containing at least 2 nuclei. Scale bar = 100 µm.

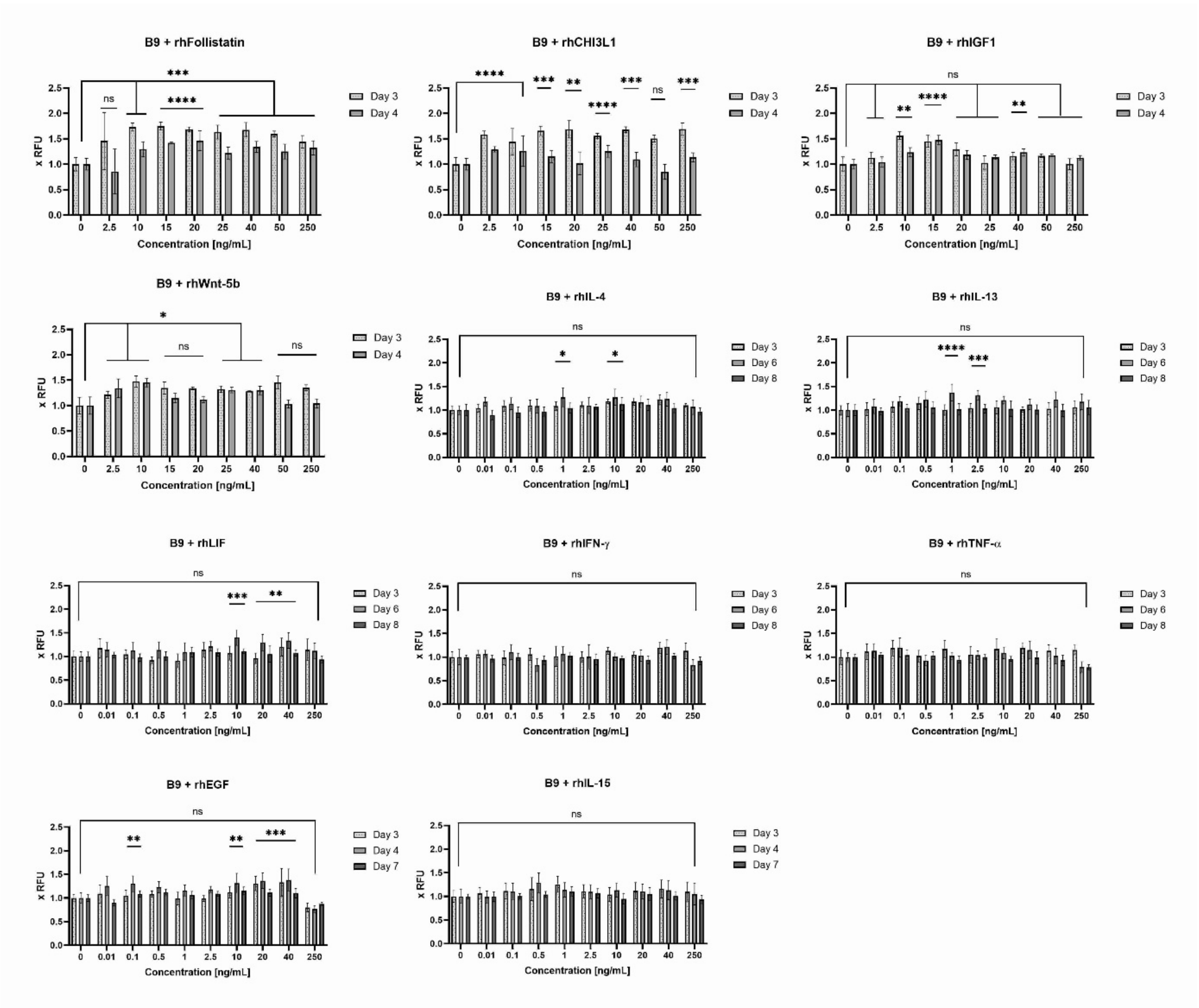

**Supplemental figure 2: Screening single medium components for induction of proliferation, Presto Blue assay.** BSCs were treated on days 1 and 3 with indicated concentrations of components, and Presto Blue assay way performed on indicated days. Obtained values were normalized to vehicle treated control (0), which contained DPBS + 0.8 mg/mL HSA (B9). n=6 biological replicates and repeated at least twice; statistical significance was calculated by one-way ANOVA combined with Tukey HSD for either day 4 or 6, comparing all samples to 100% B9, and is indicated by asterisks, which are $p < 0.05$ (*), $p < 0.01$ (**), $p < 0.001$ (***), $p < 0.0001$ (****).

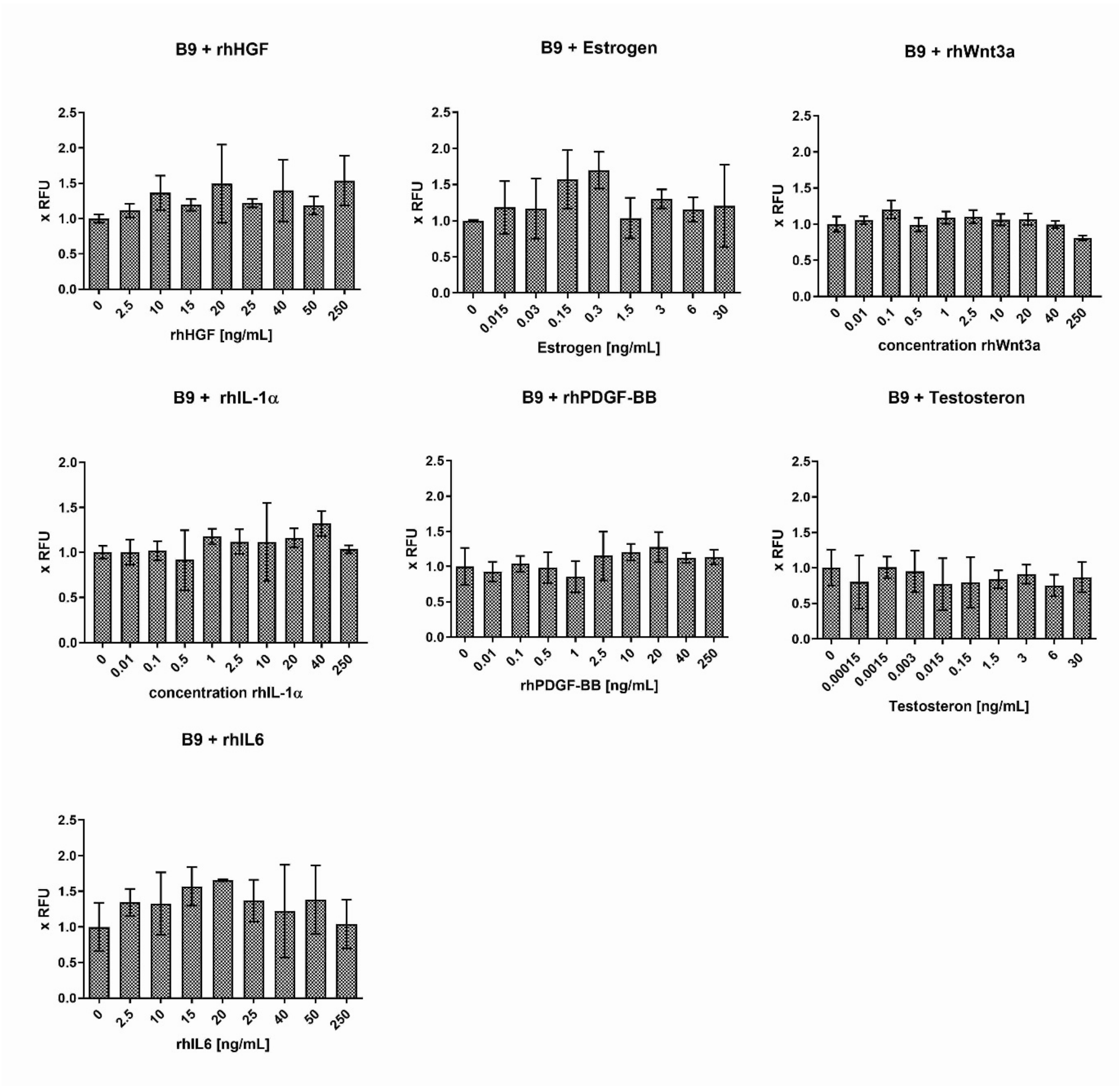

**Supplemental figure 3: Screening single medium components for induction of proliferation.** BSCs were treated on days 1 and 3 with indicated concentrations of components, and Hoechst assay way performed on day 4 (rhHGF, IL-6, Estrogen) or day 8 (rhPDGF-BB, testosterone, rhWnt3a, rhIL-1alpha). Obtained values were normalized to vehicle treated control (0), which contained DPBS + 0.8 mg/mL HSA (B9). n=3 (rhHGF, IL-6, Estrogen) and n=6 (rhPDGF-BB, testosterone, rhWnt3a, rhIL-1α) biological replicates, experiment repeated at least twice.

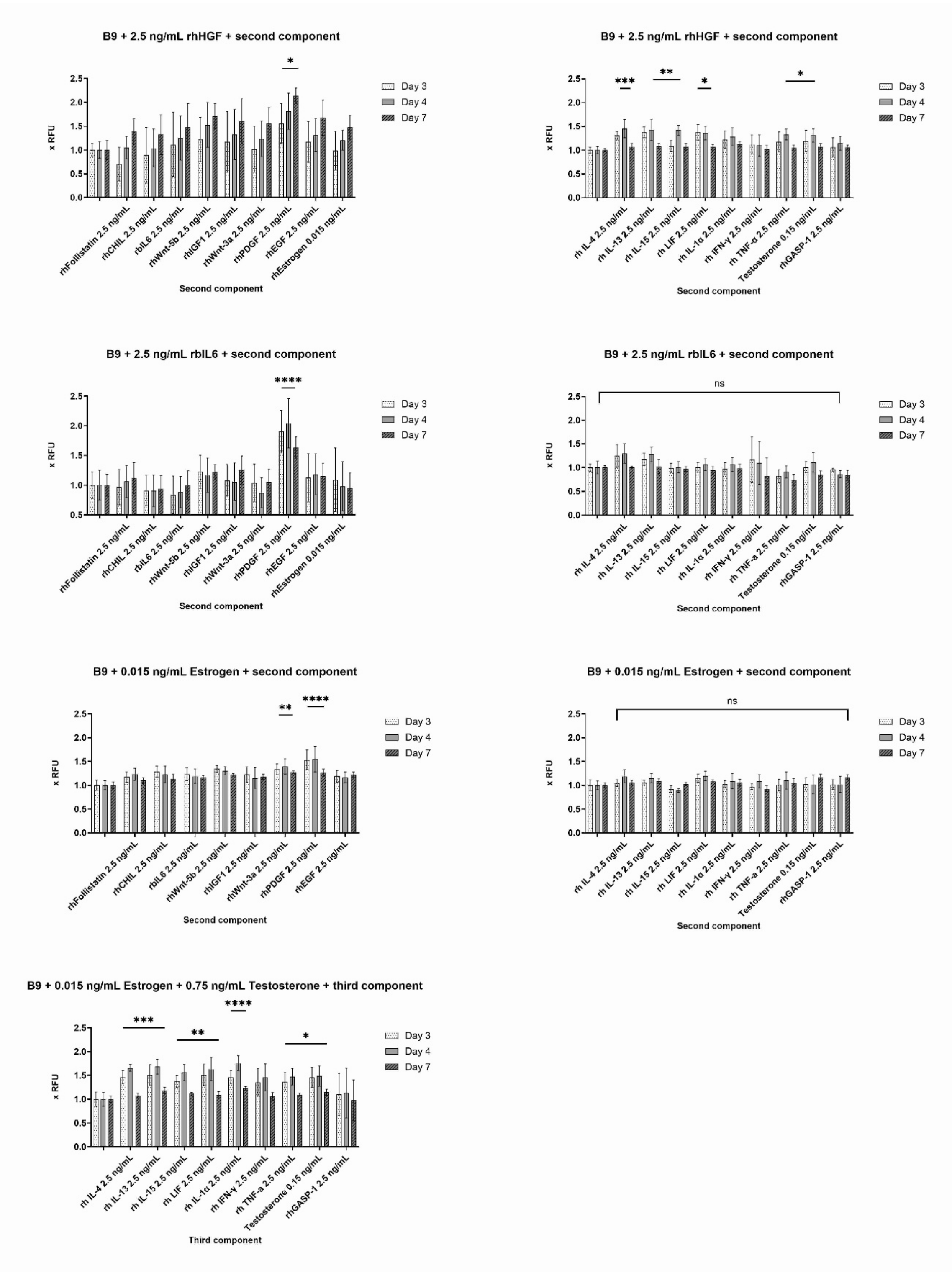

**Supplemental figure 4:** Screening combinations of medium components for induction of proliferation. BSCs were treated twice with a basic concentration of 2.5 ng/mL for protein components and 0.015 ng/mL for Estrogen and Testosterone. Presto Blue assay way performed on indicated days. Obtained values were normalized to the cells treated with only one of the hits from single component screenings (control). n=6 biological replicates and repeated at least twice; statistical significance was calculated by one-way ANOVA combined with Dunnett test for day 4, comparing all samples to B9 with indicated first component, and is indicated by asterisks, which are p < 0.05 (*), p <0.01 (**), p < 0.001 (***), p < 0.0001 (****).

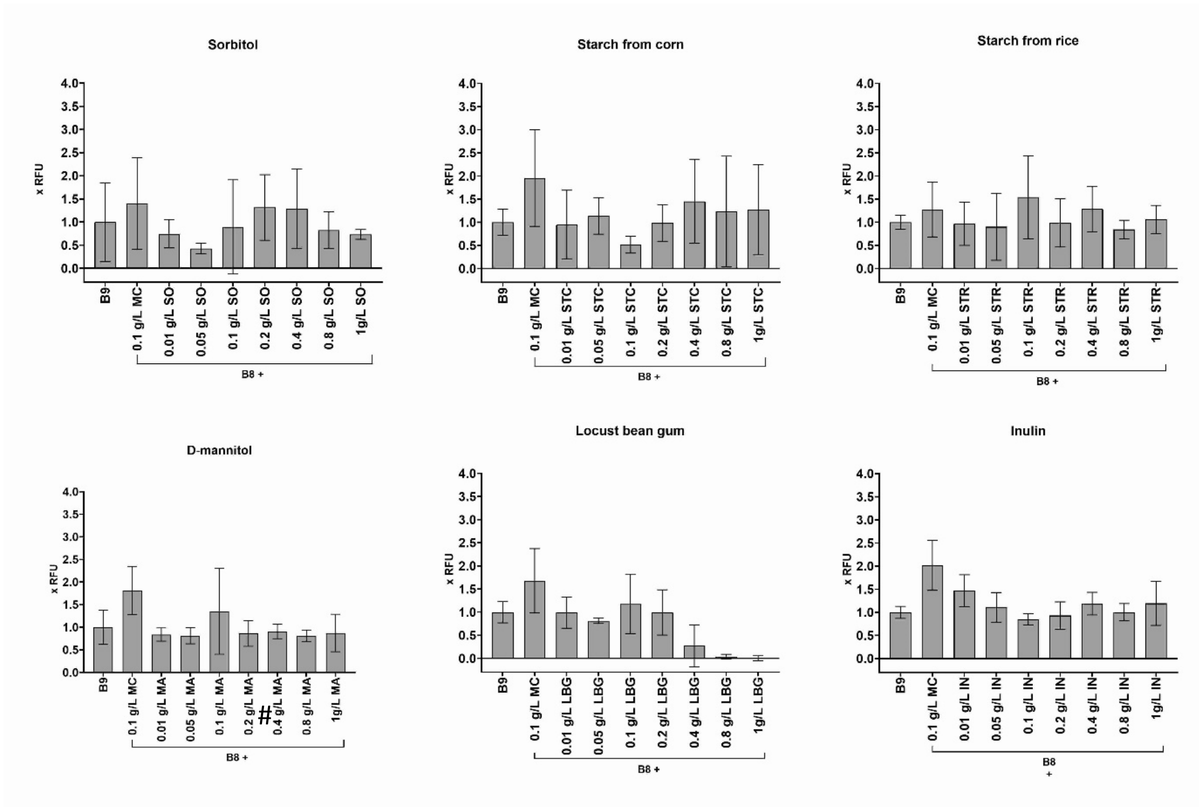

**Supplemental figure 5:** Non-specific stabilization of cultured medium components by new potential stabilizers, Hoechst assay. 2000 BSCs/cm$^2$ were seeded on day 0 in BSC-GM, and changed on day 1 to the designated medium. B8 with 0.1 g/L methyl cellulose (MC) and B9 medium were used as controls. Stabilizers were added to indicated end-concentrations with every medium exchange. Hoechst assay was performed on the last day of the experiment. Obtained values were normalized to BSCs in B9. n=6 biological replicates and repeated at least twice.

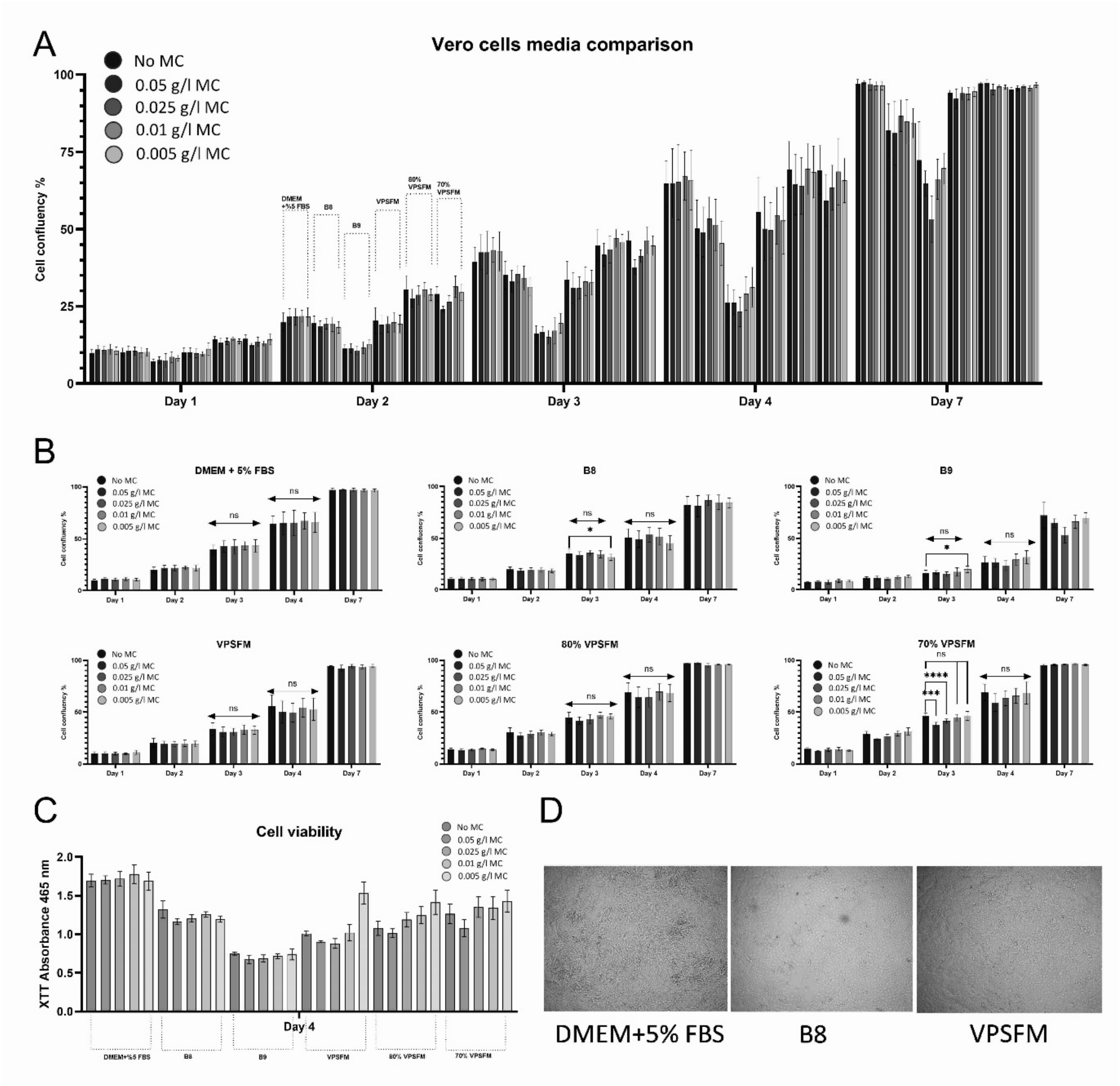

**Supplemental figure 6:** Effect of stabilizers MC and HSA on Vero cell proliferation. 10,000 cells/cm$^2$ were seeded on day 0 into designated medium. DMEM + 5% FBS and commercially available medium VPSFM were used as controls. Stabilizers were added to the indicated final concentrations at the time of seeding and medium exchange with fresh medium (4$^{th}$ day of culture). Confluence measurement was performed using live-cell imaging on TECAN device **(A-B)**, and the cell proliferation XTT assay (Roche) was performed according to the manufacturer instruction **(C)** in the designated days of the experiment. n=12 biological replicates; statistical significance was calculated by one-way ANOVA combined with the Kruskal-Wallis test for days 3 and 4, and is indicated by asterisks, which are $p < 0.05$ (*), $p < 0.01$ (**), $p < 0.001$ (***), $p < 0.0001$ (****). Microscopical observation of the cells is performed to compare the cells morphology at 7$^{th}$ day of culture **(D)**.

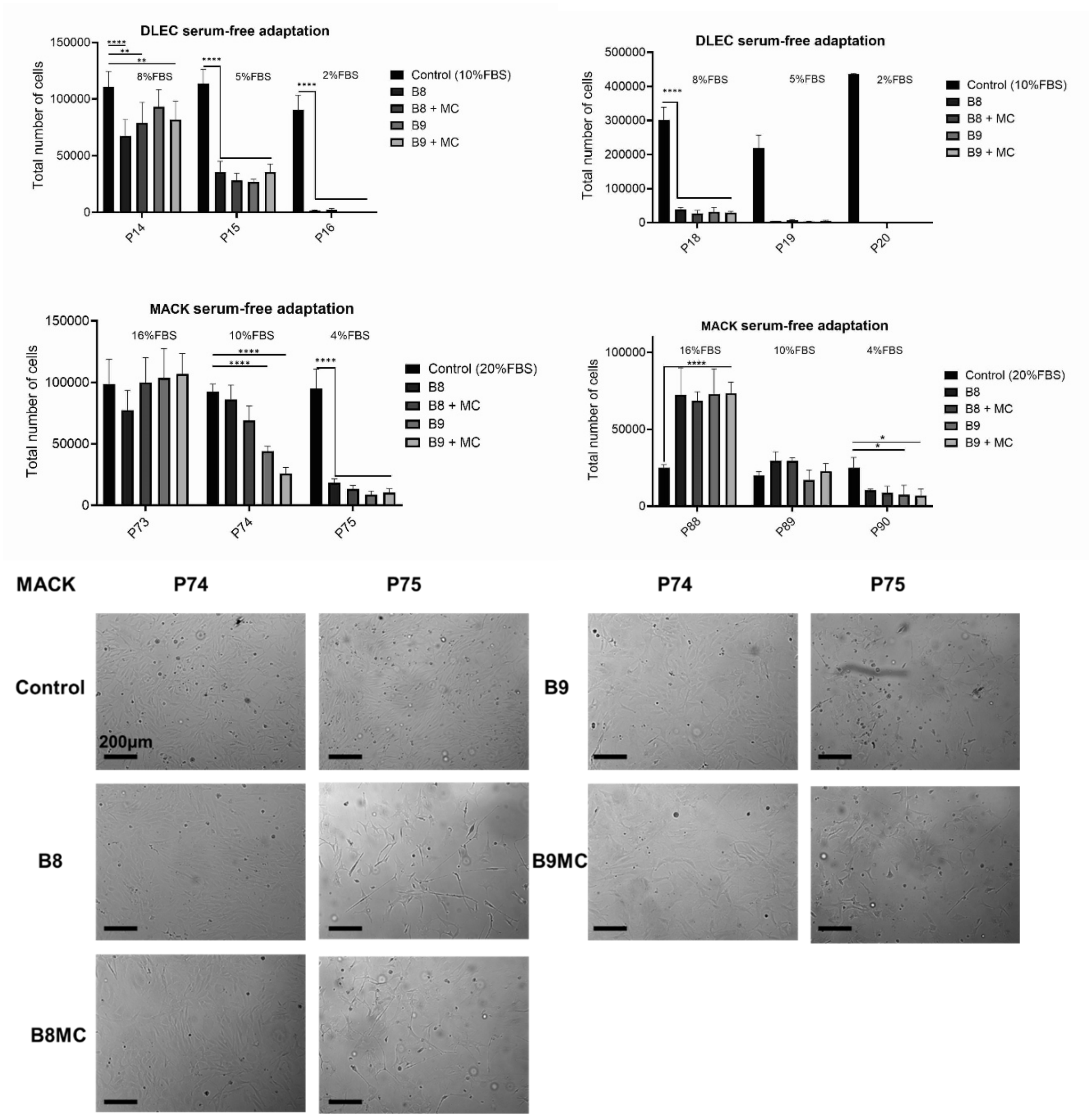

**Supplemental figure 7:** Effect of stabilizers MC and HSA on propagation of DLEC and MACK1 cells. 10000 cells/cm$^2$ were seeded on day 0 into designated medium. L15 + 10% FBS (DLEC) and L15 + 20% FBS (MACK1) were used as controls. Stabilizers were added to indicated end-concentrations with every medium exchange. Cell number was assessed using a haemocytometer and trypan-blue staining (N=2; n=8). Statistical significance was calculated by two-way ANOVA combined with Dunnett's test, and is indicated by asterisks, which are $p < 0.05$ (*), $p < 0.01$ (**), $p < 0.001$ (***), $p < 0.0001$ (****). Microscopical analysis of the cells was performed to compare the cells morphology throughout the experiment.

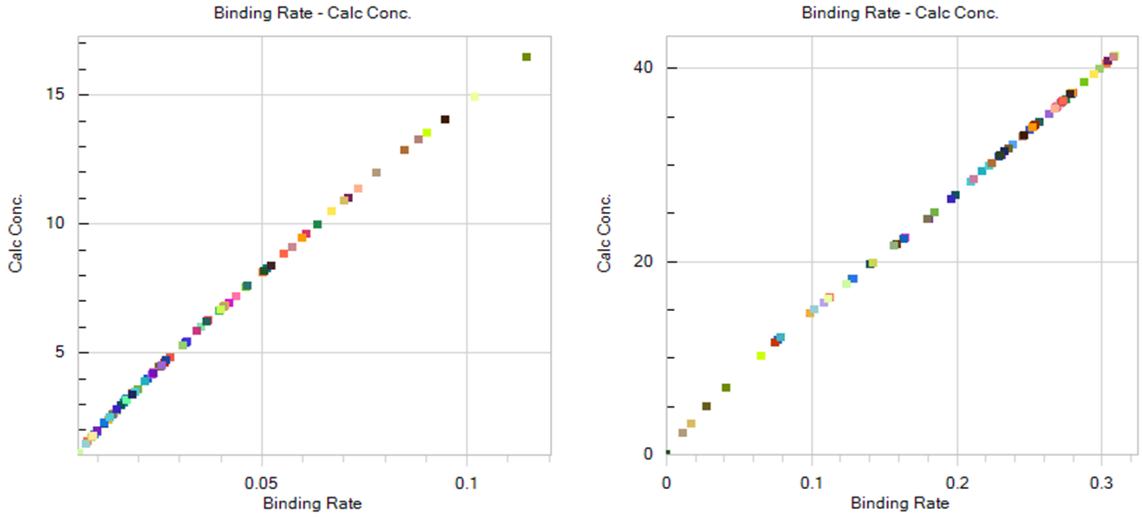

**Supplemental figure 8:** Binding rates of EPOFc in all tested samples was unaffected by the media supplementation with stabilizers, irrespective of the expression timepoint and stabilizer concentration. To assess the binding rate and concentration of the product, Octet RED96e (ForteBio) was used, sensor type - Protein A. **(A)** Samples in biological triplicates from 24h-96h, **(B)** samples in biological triplicates from 96h-192h expression in GM (CD CHO + 8mM L-Gln + 0.2%ACA), GM + 0.1g/L methyl cellulose and GM + 0.1g/L methyl cellulose + 0.8g/L HSA.

**Supplemental Table 2:** Comparison of three media regarding costs for stabilizing ECM proteins.

| **Kolkmann *et al.*[18]** | | | | |
|---|---|---|---|---|
| **Component** | **Component end concentration** | **Component bulk cost** | **Component cost Euro/L medium** | **Link** |
| ITSE Animal-Free | 1% | 488.64 € per 100 mL | 4.89 | https://invitria.com/products/itse-af-recombinant-transferrin/ |
| Glutamax | 1% | 77.00 € per 100 mL | 7.7 | https://www.thermofisher.com/order/catalog/product/35050061 |
| HSA | 5 mg/mL | 25 € per 1 g | 125 | https://www.oryzogen.net/list/6.html |
| Fibronectin | 10 µg/mL | 35 € per 1 mg | 350 | https://www.oryzogen.net/list/6.html |
| Hydrocortisone | 36 ng/mL | 305 € per 10g | 0.0011 | https://www.sigmaaldrich.com/ |
| Human IL-6 | 20 ng/mL | 2965 € per 1 mg | 59.3 | https://www.peprotech.com/de/search?q=il+6 |
| alpha linonelic acid | 1 µg/mL | 1680 € per 10 g | 0.168 | https://www.sigmaaldrich.com/AT/de/product/sigma/l2376 |
| L-acsorbate-2-phosphat | 50 µg/mL | 513 € per 100 g | 0.26 | https://www.sigmaaldrich.com/AT/de/product/sigma/49752 |
| FGF 2 | 10 ng/mL | 22 € per 1 mg | 0.22 | https://www.oryzogen.net/list/6.html |
| HGF | 5 ng/mL | 4,735.00 € per 1 mg | 23.68 | https://www.peprotech.com/en/recombinant-human-hgf-insect-derived |
| VEGF | 10 ng/mL | 150 € per 1 mg | 1.5 | https://www.oryzogen.net/list/6.html |
| IGF-1 | 100 ng/mL | 30 € per 1 mg | 3 | https://www.oryzogen.net/list/6.html |
| PDGF-BB | 10 ng/mL | 3,845.00 € per 1 mg | 38.45 | https://www.peprotech.com/en/recombinant-human-pdgf-bb |
| | | **Sum** | **614.16** | |
| | | **Sum HSA→MC+STC** | **489.21** | 0,4 g/L STC → 0.017 €/L https://www.sigmaaldrich.com/AT/de/product/sial/s4126<br>0.1125 g/L MC → 0.037 €/L https://www.sigmaaldrich.com/AT/en/product/sigma/m0512 |
| **Stout *et al.*[17]** | | | | |
| Insulin | 20 µg/mL | 41,920 € per 100 g | 8.38 | https://invitria.com/ |
| Ascorbic acid 2-phosphate | 200 µg/mL | 513 € per 100 g | 1.03 | https://www.sigmaaldrich.com/AT/de/product/sigma/49752 |
| Transferrin | 20 µg/mL | 70 € per 1 g | 1.4 | https://www.oryzogen.net/list/6.html |
| Sodium selenite | 20 ng/mL | 240 € per 100 g | 0.000048 | https://www.sigmaaldrich.com/AT/de/product/sigma/s5261 |
| FGF2-G3 | 40 ng/mL | 22€ per 1 mg | 0.22 | https://www.oryzogen.net/list/6.html |
| TGFβ3 | 0.1 ng/mL | 1,235 € per 100 µg | 1.24 | Recombinant human TGF beta 3 protein (Active) (ab269208) \| Abcam |
| NRG1 | 0.1 ng/mL | 1,484.00 € per 1 mg | 0.15 | https://www.peprotech.com/en/recombinant-human-heregulin-1 |
| Sodium bicarbonate | 2438 µg/ml | 59.30 € per 500 g | 0.29 | https://www.sigmaaldrich.com/AT/de/product/sigma/s5761 |
| Albumin | 800mg/L | 25 € per 1 g | 20 | https://www.oryzogen.net/list/6.html |
| | | **Sum** | **32.70** | |
| | | **Sum HSA→MC+STC, % of total** | **12.75** | 0,4 g/L STC → 0.017 €/L https://www.sigmaaldrich.com/AT/de/product/sial/s4126<br>0.1125 g/L MC → 0.037 €/L https://www.sigmaaldrich.com/AT/en/product/sigma/m0512 |
| **Skrivergaard *et al.*[19]** | | | | |
| FGF-2 | 2 ng/mL | 800 € per 1 mg | 32 | https://www.peprotech.com/en/recombinant-human-fgf-basic-154-aa |
| Fetuin | 600 µg/mL | 69.20 € per 100 mg | 415.2 | https://www.sigmaaldrich.com/ |
| BSA | 75 µg/mL | 154 € per 100 mL 7.5% in DPBS (x100) | 15.4 | https://www.sigmaaldrich.com/AT/de/product/sigma/a8412 |
| ITSE Animal-Free | 1% | 488.64 € per 100 mL | 4.89 | https://www.thermofisher.com/order/catalog/product/51500056 |
| PDGF | 5 ng/ml | 3,845.00 € per 1 mg | 38.45 | https://www.peprotech.com/en/recombinant-human-pdgf-bb |
| HGF | 20 ng/ml | 4,735.00 € per 1 mg | 23.68 | https://www.peprotech.com/en/recombinant-human-hgf-insect-derived |
| | | **Sum** | **529.62** | |
| | | **Sum BSA→MC+STC** | **514.27** | |
| | | **Sum BSA+Fetuin→MC+STC** | **99.07** | 0,4 g/L STC → 0.017 €/L https://www.sigmaaldrich.com/AT/de/product/sial/s4126<br>0.1125 g/L MC → 0.037 €/L https://www.sigmaaldrich.com/AT/en/product/sigma/m0512 |
